# Molecular mechanisms involved in Atlantic halibut (*Hippoglossus hippoglossus*) egg quality: impairments at transcription and protein folding levels induce inefficient protein and energy homeostasis during early development

**DOI:** 10.1101/2022.02.01.478612

**Authors:** Ozlem Yilmaz, Anders Mangor Jensen, Torstein Harboe, Margareth Møgster, Ragnfrid Mangor Jensen, Olav Mjaavatten, Even Birkeland, Endy Spriet, Linda Sandven, Tomasz Furmanek, Frode S. Berven, Anna Wargelius, Birgitta Norberg

## Abstract

**Background:** Reproductive success and normal development in all animals are dependent on egg quality and developmental competence of the produced embryo. This study employed tandem mass tags labeling based liquid chromatography tandem mass spectrometry for egg proteomic profiling to investigate differences in the global proteome of good versus poor quality Atlantic halibut eggs at 1-cell stage post fertilization.

**Results:** A total of 115 proteins were found to be differentially abundant between good and poor quality eggs. Frequency distribution of these proteins revealed higher protein folding activity in good quality eggs in comparison to higher transcription and protein degradation activities in poor quality eggs (*p* < 0.05). Poor quality halibut eggs were significantly enriched with additional proteins related to mitochondrial structure and biogenesis (*p* < 0.05). The differential abundance of a selection of proteins was first confirmed at gene expression level using a transcriptomic approach followed by a targeted proteomic approach (parallel reaction monitoring based mass spectrometry) in biological samples obtained from two consecutive reproductive seasons. The findings of global proteome profiling, together with the validation of differential abundance of targeted proteins and their related genes, suggest impairments in protein and energy homeostasis which might be related to unfolded protein response and mitochondrial stress in poor quality eggs. Additional transmission electron microscopy studies were taken to assess potential differences in abundance and morphological integrity of mitochondria between good and poor quality eggs. Observations reveal poor quality eggs to contain significantly higher number of mitochondria with higher number of cristae. These mitochondria, however, are significantly smaller and have a more irregular shape than those found in high-quality eggs. Therewithal difference in mtDNA levels represented by *mt-nd5* and *mt-atp6* genomic DNA abundance in this study, were found to be not statistically significant (*p* > 0.05) between good and bad quality eggs at both 1 hpf and 24 hpf stages.

**Conclusion:** Overall evidence from this study indicate that poor quality eggs undergo impairments at both transcription and translation level leading to endoplasmic reticulum and mitochondrial deficiencies. Additional research may be required to expediate the details and the potential of these impairments occurring in different species. Nonetheless, this study will pave the way for future research and will help in acceleration of recent advances in the field of embryonic developmental competence of living organisms.

## BACKGROUND

Egg quality is of pivotal importance in biomedicine, agriculture, ecology and environmental science because of its tremendous influence on reproductive success or failure in all animals. Poor egg quality remains a serious problem of largely variable cause(s) in human reproductive medicine (1,2) (Tarin et al., 2014, Keefe et al., 2014) and livestock production (3–5) (Kjorsvik et al., 1990, Bobe and Labbe, 2010, Migaud et al., 2013). Maternal factors, primarily mRNA and proteins deposited in the egg during ovarian expansion and maturation, are among the key influences on fertility and early embryogenesis. Recent research focus has been increasingly devoted to investigating the motherlode of maternal mRNA and proteins deposited in the egg for clues to the origin of egg quality problem and possible solutions (6–13) (Aegerter et al., 2005, Bonnet et al. 2007, Mommens et al. 2014, Chapman et al. 2014, Sullivan et al., 2015, Zarski et al., 2017, Cheung et al., 2019, Ma et al., 2019). In vertebrates the maternal RNA stockpile and proteins drive the early embryonic development until activation of the zygotic genome around mid-blastula stage (14,15) (Tadros and Lipshitz, 2009, Jukam et al., 2018). Differential abundance of maternal transcripts may be indicators of quality in fish eggs (9) (Chapman et al., 2014), however, certain molecular changes such as the modification of proteins after their uptake into growing oocytes play crucial roles in many aspects of early development. These roles are not possible to determine using transcriptomic technologies and need application of proteomics, an approach representing the dynamic transfer of genetic information into the actual effector molecules in the cell, for elucidation of ongoing cellular events prior to zygotic genome activation. Despite the restricted consistency between transcript and product protein abundances (16) (Groh et al., 2011) validation of proteomic changes via transcriptomic approaches may also be applied in steady-state cellular mechanisms at early stages of development.

Proteomic profiling has been widely employed to study the cell biology of oocytes in many species, including humans, mice, pigs, fish and insects (17) (Chapovetsky et al., 2007), but it has been less than two decades since it has been considered as a useful and practical tool to study fish egg quality, i.e. rainbow trout (Oncorhynchus mykiss) (18) (Rime et al., 2004), European sea bass (Dicentrarchus labrax) (19) (Crespel et al., 2008), Eurasian perch (Perca fluviatilis) (20) (Castets et al., 2012), and hapuku (Polyprion oxygeneios) (21) (Kohn et al., 2015). Our recent research comparing the global proteomes of different quality eggs from zebrafish revealed a number of proteins as potential markers, but more importantly, several molecular mechanisms and related physiological processes to be associated with egg quality in this species (22) (Yilmaz et al., 2017). In a most up to date study, the consecutive changes in global proteome of 1-cell-stage egg after invalidation of certain types of Vtgs (*vtg1*, *4* and *5*; *vtg1*-KO and *vtg3*; *vtg3-KO*) were investigated using CRISPR/Cas9 genome editing technology (23,24) (Yilmaz et al., 2019, Yilmaz et al., 2021). The collective results of these studies delivered a clear portrait of the impaired molecular mechanisms that impacts egg and offspring developmental competence with striking similarities between *vtg*-KO and poor quality egg proteome profiles in zebrafish.

Despite species specific differences in physiological aspects of early development, the evolutionary conserved stereotypical procedure of cellular events, led us to investigate whether these findings are common with marine species of aquaculture interest. Atlantic halibut (*Hippoglossus hippoglossus*) a highly prized species in global fish markets with decreasing landings in capture fisheries and increasing demand to its production is considered as a representative of such species. Notwithstanding the progress in research and cultivation efforts that has been made recently, several persisting bottlenecks (i.e. the unsteady supply of high quality eggs and fry) still restrain expansion of sustainable commercial production of Atlantic halibut. As a batch spawner releasing up to 10 batches of eggs with highly variable quality at 2-3 days intervals during each reproductive cycle halibut is a perfect candidate to study egg quality related mechanisms.

Therefore, the objectives of this study were 1) to reveal the proteomic profiles of good versus poor quality eggs, 2) to identify proteins that can serve as egg quality markers, and 3) to discover molecular mechanisms determining egg quality using a combination of the most advanced proteomics approaches such as tandem mass tags (TMT) labeling and parallel reaction monitoring (PRM) based liquid chromatography tandem mass spectrometry (LC-MS/MS) practices. Discoveries of such mechanisms in poor quality eggs will spur development of practical strategies to determine and eliminate the potential causes leading to egg quality problems in Atlantic halibut and other farmed fishes, thereby contributing significantly to development of effective strategies for improving breeding practices and sustainable growth of Norwegian and global aquaculture. The findings of this study will also contribute considerably to recent advancements in reproductive biology of other living organisms, such as animals and humans, that share common properties of existence and cellular events.

## RESULTS

The egg batches from halibut females employed in this study showed high variation in fecundity, buoyancy, fertilization, and normal cell division ratios with no obvious link to the embryo survival ratio prior hatching at 12 days post fertilization (dpf) (**Table S1, Fig S1**). As a result, despite the high percentage of fertilization and embryo going through normal cell division processes, poor quality eggs exhibited low embryo survival rates. Based on our overall experience in hatchery practices, the cumulative percent of embryo survival for all batches stabilized prior hatching (by 12 dpf). Therefore, embryo survival at this stage was utilized as the measure of egg quality in this study. The actual survival rates in the overall samples inventory ranged from 93 % for good quality eggs and 25 % for the poor quality eggs. Due to yearly changes in this index the survival difference window between good and poor quality egg batches were not possible to standardize. Therefore, egg batches with embryo survival rate of ≥ 76 were considered to be of good quality and those spawns with ≤ 62 embryo survival were considered to be of poor quality in 2019. This ratio was ≥ 76 % and ≥ 70 % for good quality egg batches, and ≤ 71 and ≤ 55 for poor quality egg batches in the years 2020 and 2021, respectively.

### TMT labeling based LC-MS/MS

A total of 1619 out of 1886 identified proteins were considered as valid based on filtering to be present in at least four biological samples. A total of 115 of valid proteins were found to be differentially abundant between good and poor quality eggs (Independent samples t-test, *p* < 0.05 followed by Benjamini Hochberg correction for multiple testing, *p* < 0.05). Detailed information on these proteins is given in **Table S2**. In this study, proteins with higher abundance in good quality eggs are indicated as down-regulated in poor quality eggs (N = 64), and those with higher abundance in poor quality eggs are indicated as up-regulated in poor quality eggs (N = 51). **Fig 1A** demonstrates hierarchical clustering of these proteins based on *p* values obtained from Student’s t-test, *p* < 0.05 followed by Benjamini Hochberg correction for multiple testing, *p* < 0.05. A heatmap representation of differentially regulated proteins’ clustering based on their abundance between good and poor quality eggs is given in **Fig 1B**.

**Fig 1.**
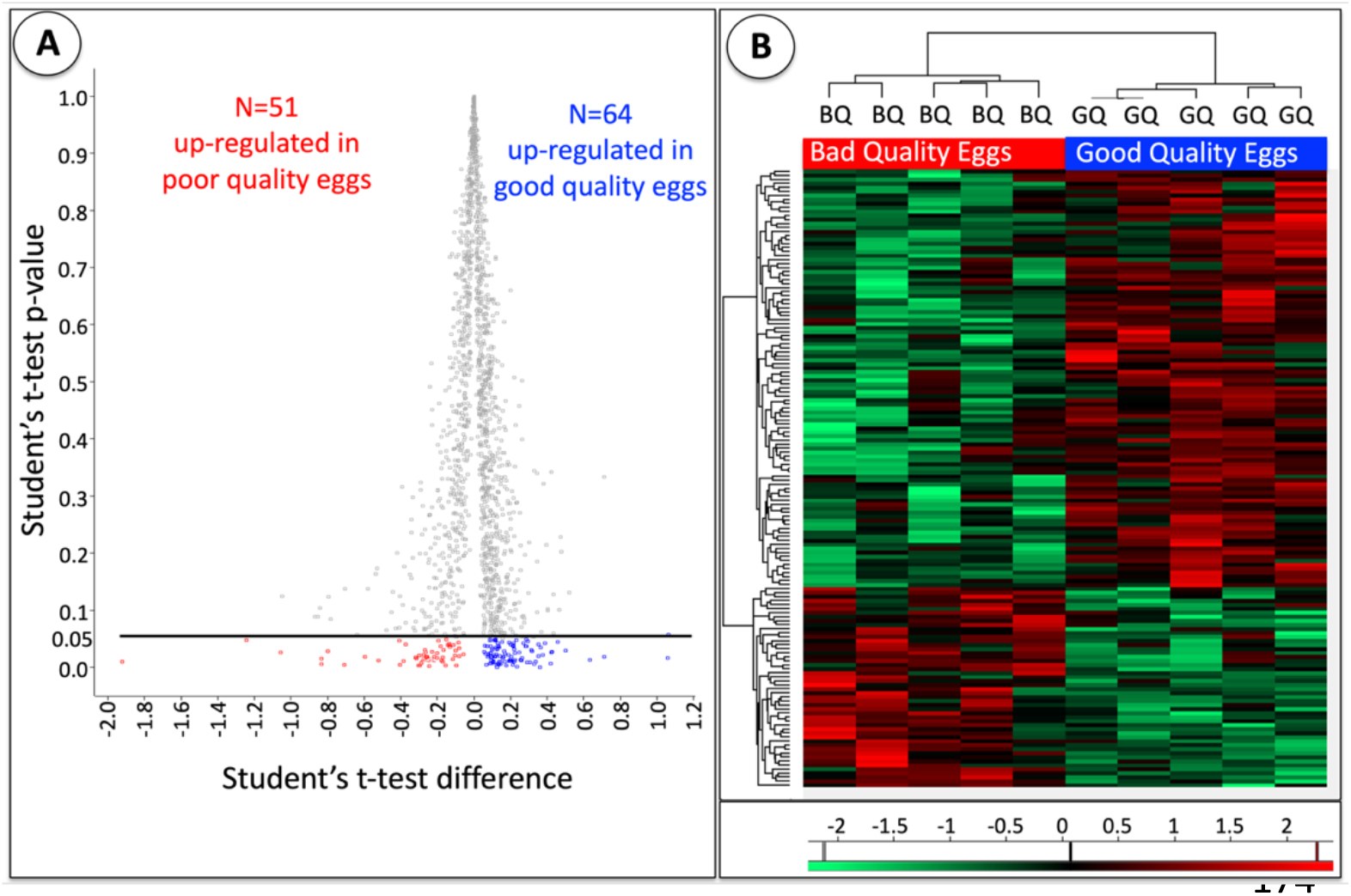
Differentially abundant proteins between good and poor quality halibut eggs. **Panel A.** Representation of differential abundance for 115 proteins detected by TMT labeling based LC-MS/MS based on their Student’s t-test significance value. Y axis indicates *p* values while X axis represents test differences. Proteins up-regulated in poor quality eggs (N = 51) are indicated in red while those which were up-regulated in good quality eggs (therefore down-regulated in poor quality eggs, N = 64)) are indicated in blue. A black horizontal line above red and blue markers representing the separation of differentially abundant proteins retained after the *p* < 0.05 cut off value. A complete list of these proteins along with detailed information on their NCBI gene IDs, NCBI accession numbers, associated protein names from human database, protein full names, functional categories (according to **Fig 2**), significance of differences in abundance (Independent t-test *p* < 0.05 followed by Benjamini Hochberg correction for multiple tests *p* < 0.05), relative abundance ratios (GQ/BQ and BQ/GQ, respectively), and regulation tendencies (BQ-upregulated or BQ-downregulated) are given in **Table S2**. **Panel B.** A heatmap clustering of differentially abundant proteins between good and poor quality egg groups.

Frequency distribution of differentially abundant proteins among different thirteen arbitrarily chosen functional categories that would account for > 90 % of the proteins is given in **Fig 2**. Accordingly, proteins which were down-regulated in poor quality eggs (N = 64) (**Fig 2 Left panel; Good Quality Eggs**) were mainly related to cell cycle, division, growth and fate (26 %), protein folding (14 %), energy metabolism (12 %), translation (11 %), protein transport (8 %), lipid metabolism (8 %) with the remaining categorized proteins being related to protein degradation and synthesis inhibition (5 %), transcription (5 %), mitochondrial biogenesis (5 %), metabolism of cofactors and vitamins (3 %), Redox/detox related (1%). Two percent of proteins which were down-regulated in poor quality eggs were placed in the category of ‘others’. Proteins which were up-regulated in poor quality eggs (N = 51) (**Fig 2 Right panel; Poor Quality Eggs**) were mainly related to cell cycle, division, growth and fate (19 %), protein degradation and synthesis inhibition (18 %), mitochondrial biogenesis (17 %), transcription (16 %), energy metabolism (8 %), protein transport (8 %) with the remaining categorized proteins being related to lipid metabolism (4 %), protein folding (2 %), translation (2 %), Redox/detox related (2), immune response related (2%). Two percent of proteins which were up-regulated in poor quality eggs were placed in the category of ‘others’. The distribution of these differentially regulated proteins among functional categories significantly differed between egg quality groups (χ^2^ *p* < 0.05). Accordingly, good quality eggs seem to contain significantly higher number of proteins related to protein folding (14 %), while poor quality eggs contain significantly higher number of proteins related to transcription (16 %), protein degradation and synthesis inhibition (18 %), and mitochondrial biogenesis (17 %) (**Fig 2**).

**Fig 2.**
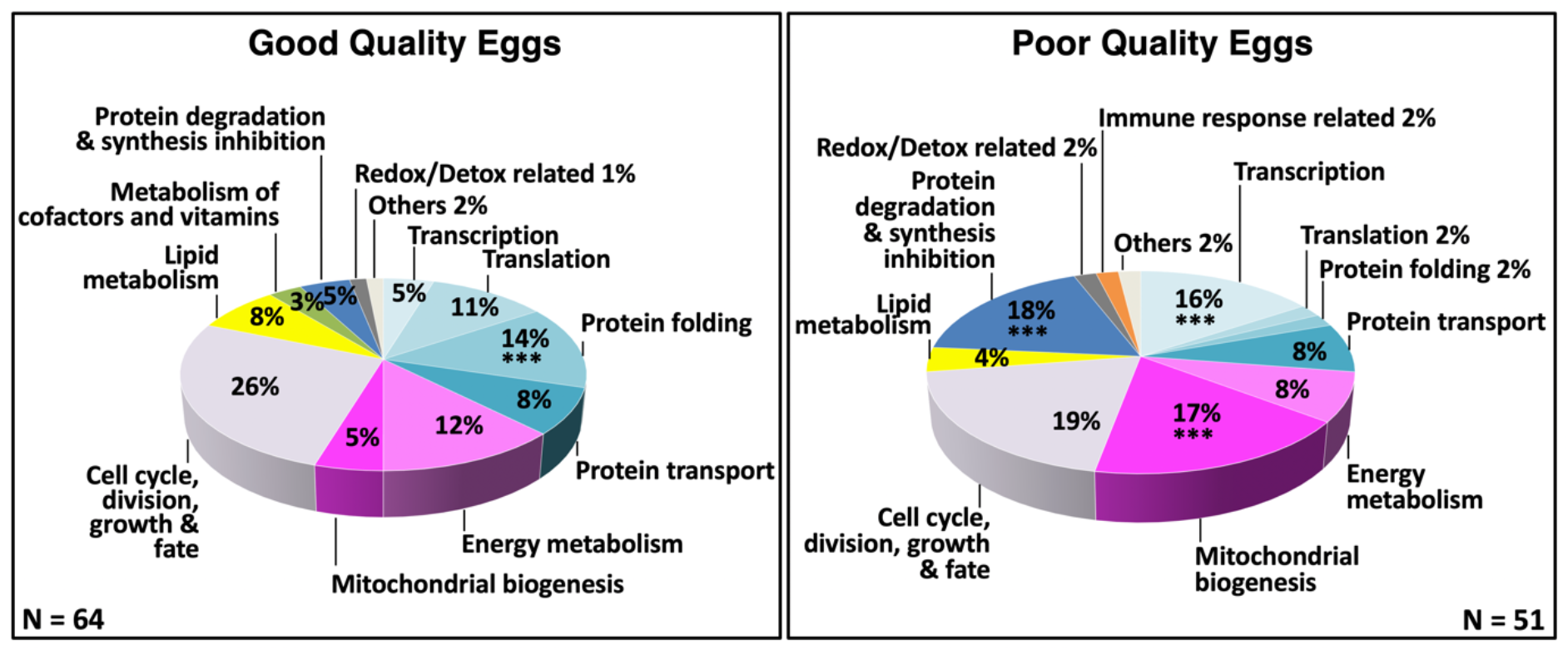
Distribution of differentially abundant proteins among functional categories. **Panel A**. Proteins up-regulated in good quality eggs (N = 64), therefore down-regulated in poor quality eggs. **Panel B.** Proteins up-regulated in poor quality eggs (N = 51). The overall distribution of differentially regulated proteins among the functional categories significantly differed between good and poor quality eggs (χ^2^, *p* < 0.05). Asterisks indicate significant differences between different groups in the proportion of differentially regulated proteins within a functional category (χ^2^, *p* < 0.05). The corresponding NCBI gene IDs, NCBI accession numbers, associated protein names from human database, protein full names, functional categories (shown above), significance of differences in abundance (Independent t-test *p* < 0.05 followed by Benjamini Hochberg correction for multiple tests *p* < 0.05), relative abundance ratios (GQ/BQ and BQ/GQ, respectively), and regulation tendencies (BQ-upregulated or BQ-downregulated) are given in **Table S2.**

Gene ontology (GO) enrichment analysis based on overrepresentation test (*p* < 0.05), with the human database being used as a reference, revealed significant biological processes, molecular functions and cellular components which are in close relation with frequency distribution analysis findings. Respectively, biological processes which were overrepresented by proteins down-regulated in poor quality eggs were as follows; protein folding, small molecule catabolic process, ribonucleoprotein (RNP) complex biogenesis, RNP complex subunit organization, cofactor biosynthetic process, coenzyme metabolic process, organophosphate (OP) catabolic process, nuclear transport, nucleobase-containing small molecule biosynthetic process, and RNA catabolic process. Molecular functions which were mostly overrepresented by poor quality down-regulated proteins were related to isomerase activity, oxidoreductase activity (acting on the aldehyde or oxo group of donors), oxidoreductase activity (acting on the CH-CH group of donors), oxidoreductase activity (acting on a sulfur group of donors), RNP complex binding, kinesin binding, translation factor activity (RNA binding), snRNA binding, mRNA binding, and Ran GTPase binding. Cellular components overrepresented by these proteins were RNP complex, Sm-like protein family complex, mitochondrion, cytoplasmic region, mitochondrial part, cytoplasmic RNP granule, P-body, mitochondrial matrix, neuron projection cytoplasm, and tertiary granule lumen. KEGG pathways that were significantly overrepresented by the same set of proteins were RNA degradation, metabolic pathways, fatty acid degradation, valine, leucine and isoleucine degradation, glycolysis/gluconeogenesis, necroptosis, propanoate metabolism, glycine, serine and threonine metabolism, tryptophane metabolism, and ferroptosis (**Fig 3A, 3B, 3C and 3D Left panels**).

**Fig 3.**
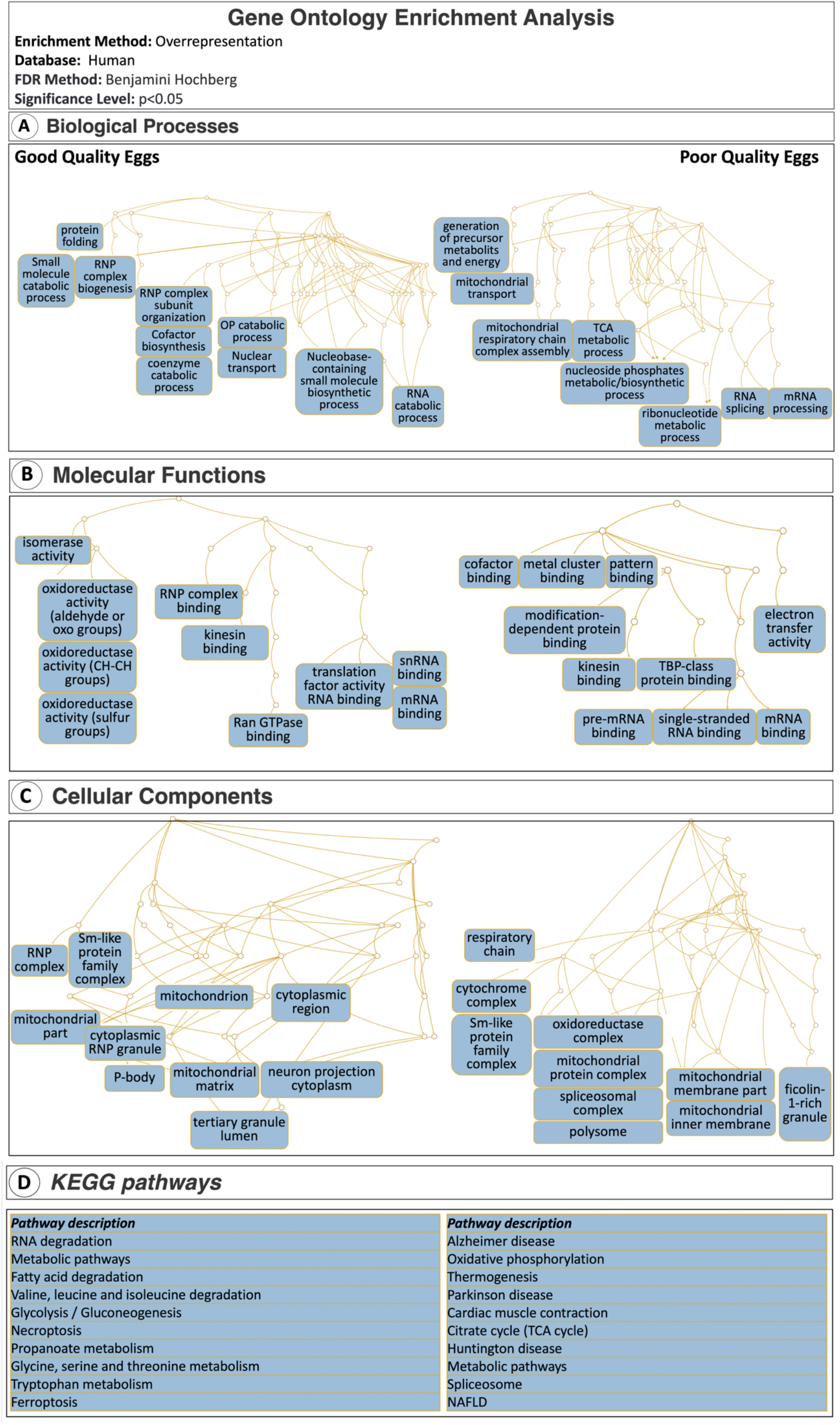
Gene ontology overrepresentation-based enrichment analyses for differentially abundant proteins. **Panel A.** Biological processes significantly enriched in good quality eggs (Left) versus in poor quality eggs (Right). **Panel B.** Molecular functions significantly enriched in good quality eggs (Left) versus in poor quality eggs (Right). **Panel C.** Cellular components significantly enriched in good quality eggs (Left) versus in poor quality eggs (Right). **Panel D.** KEGG pathways significantly enriched in good quality eggs (Left) versus in poor quality eggs (Right). A total of N = 51 and N=64 proteins which were up- and down-regulated in poor quality eggs were mapped against human database for enrichment analyses using the overrepresentation method at *p* < 0.05 followed by Benjamini Hochberg correction for multiple testing (*p* < 0.05).

In contrast, biological processes which were overrepresented by proteins up-regulated in poor quality eggs were as follows; generation of precursor metabolites and energy, mitochondrial transport, mitochondrial respiratory chain complex assembly, tricarboxylic acid metabolic process, nucleoside phosphates metabolic/biosynthetic process, ribonucleotide metabolic process, RNA splicing, and mRNA processing. Molecular functions which were mostly overrepresented by poor quality down-regulated proteins were related to cofactor binding, metal cluster binding, pattern binding, modification-dependent protein binding, kinesin binding, TBP-class protein binding, electron transfer activity, pre-mRNA binding, single-stranded RNA binding, and mRNA binding. Cellular components overrepresented by these proteins were respiratory chain, cytochrome complex, Sm-like protein family complex, oxidoreductase complex, mitochondrial protein complex, spliceosomal complex, polysome, mitochondrial membrane part, mitochondrial inner membrane and ficolin-1-rich granule. KEGG pathways which were significantly overrepresented by the same set of proteins were Alzheimer disease, oxidative phosphorylation, thermogenesis, Parkinson’s disease, cardiac muscle contraction, citrate cycle (TCA cycle), Huntington disease, metabolic pathways, spliceosome, and non-alcoholic fatty liver disease (NAFLD) (**Fig 3A, 3B, 3C and 3D Right panels**).

When the 115 differentially regulated proteins with significant differences in abundance between good and poor quality eggs were submitted separately (down-regulated in BQ; N = 64, up-regulated in BQ; N = 51) to a functional protein association networks analysis using the Search Tool for the Retrieval of Interacting Genes/Proteins (STRING) and the human protein database, they resolved into networks with significantly and substantially greater numbers of known and predicted interactions between proteins than would be expected of the same size lists of proteins randomly chosen from the human database (**Fig 4**). The subnetwork formed by proteins down-regulated in poor quality eggs is made up of three major interrelated clusters mainly related to cytoskeletal regulation, energy and protein homeostasis (**Fig 4 Left panel**). A subcluster to the far left includes proteins involved in cytoskeletal organization such as Adenylyl cyclase-associated protein 1 (CAP1), Actin beta (ACTB), Tubulin alpha 4a (TUBA4A), Kinesin family member 1B (KIF1B), Voltage dependent anion channel 1 (VDAC1), Deoxyuridine 5’-triphosphate nucleotidohydrolase, mitochondrial (DUT), Adenosylhomocysteinase like 1 (AHCYL1) and in energy production and homeostasis such as Creatine kinase (M-type) (CKM), Phosphoglycerate mutase 1 (PGAM1), Enolase (ENO1). Other proteins forming this cluster are the Complement component 1 Q subcomponent-binding protein, mitochondrial (C1QBP), and Prohibitin (PHB) which are related to mitochondrial structure and the Superoxide dismutase 1 (SOD1) which is related to redox/detox activities.

**Fig 4.**
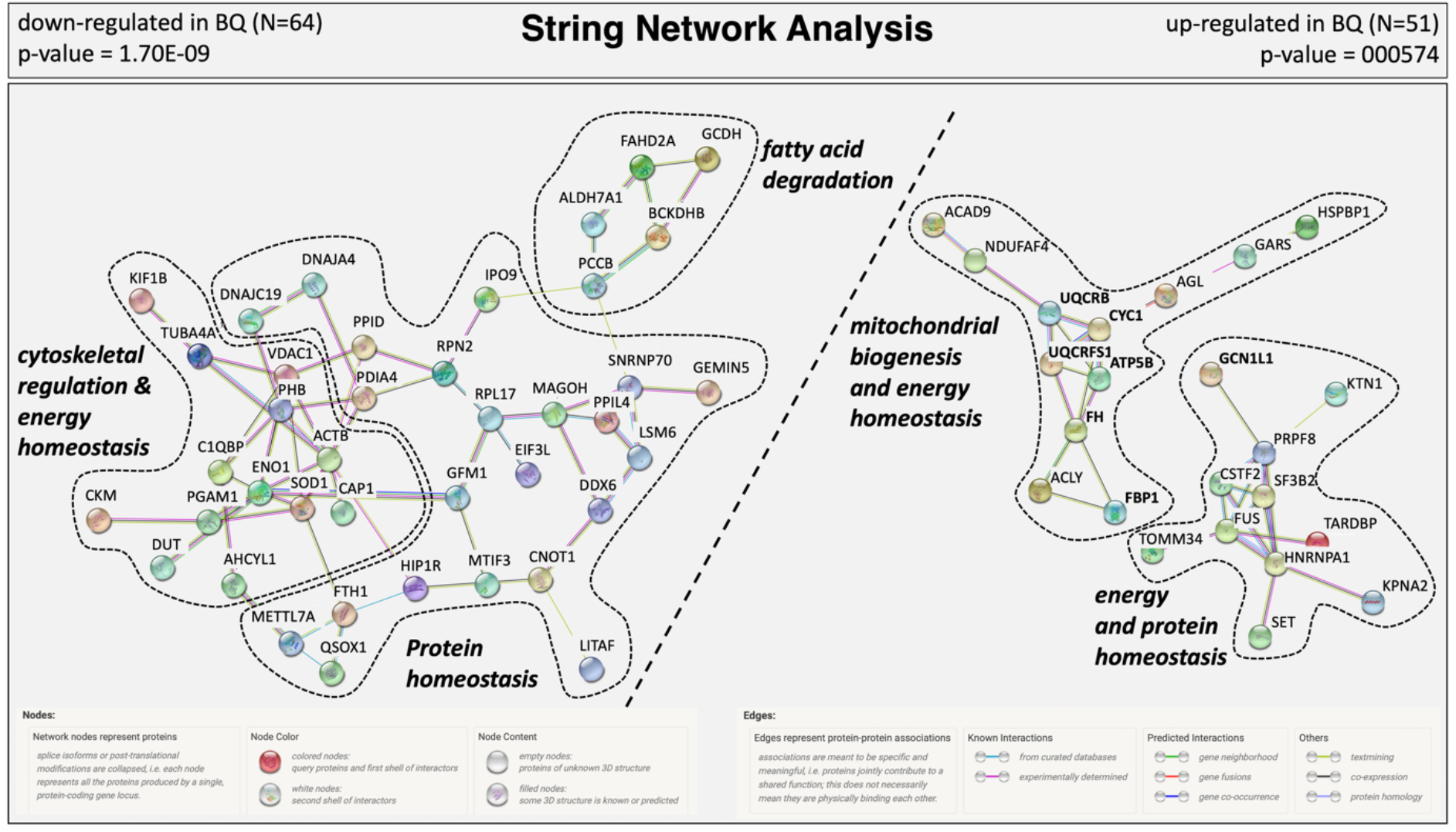
STRING Network Analysis of the differentially abundant proteins. Protein-protein interactions network clusters are given for N = 64 proteins which were down-regulated in poor quality (BQ) eggs and N = 51 proteins which were up-regulated in BQ eggs. The subnetworks formed by proteins down-regulated in BQ eggs are shown to the upper left above the diagonal dashed line, and the subnetworks formed by proteins up-regulated in BQ eggs are shown to the lower right below the diagonal dashed line. Where possible, dashed lines encircle clusters of interacting proteins involved in physiological processes distinct from other such clusters. Each network node (sphere) represents all proteins produced by a single, protein-coding gene locus (splice isoforms or post-translational modifications collapsed). Only nodes representing query proteins are shown. Nodes are named for the human proteins to which spectra were mapped; for full protein names, see **Tables S2**. Edges (colored lines) represent protein-protein associations meant to be specific and meaningful, e.g. proteins jointly contribute to a shared function but do not necessarily physically interact. Model statistics are presented at the top left and at the top right of each panel for proteins down- and up-regulated in BQ eggs, respectively. Explanation of edge colors is given below panels.

The cluster to the right of the revealed network covers proteins related to mRNA biogenesis and transcription (i.e. LSM6 homolog, U6 small nuclear RNA and mRNA degradation associated (LSM6), ATP-dependent RNA helicase DDX6 (DDX6), Small nuclear ribonucleoprotein U1 subunit 70 (SNRNP70), Mago homolog, exon junction complex subunit (MAGOH)), protein translation (i.e. Gem-associated protein 5 (GEMIN5), Eukaryotic translation initiation factor 3 subunit L (EIF3L), Ribosomal protein L17 (RPL17), G elongation factor mitochondrial 1 (GFM1), and the Translation initiation factor IF-3, mitochondrial (MTIF3)), protein folding (i.e. Dolichyl-diphosphooligosaccharide-protein glycosyltransferase subunit 2 (RPN2), Methyltransferase like 7A (METTL7A), Peptidylprolyl isomerase like 4 (PPIL4), DnaJ heat shock protein family (Hsp40) member A4 (DNAJA4), DnaJ heat shock protein family (Hsp40) member C19 (DNAJC19), Peptidylprolyl isomerase D (PPID), Protein disulfide isomerase family A member 4 (PDIA4), Quiescin sulfhydryl oxidase 1 (QSOX1), and protein transport (i.e. Importin (IPO9) and the Huntingtin-interacting protein 1-related protein (HIP1R)). Three other proteins covered by this subcluster are CCR4-NOT transcription complex subunit 1 (CNOT1) a transcription suppressor in DNA damage, Lipopolysaccharide-induced tumor necrosis factor-alpha factor homolog (LITAF) which targets proteins for lysosomal degradation, and Ferritin heavy chain 1 (FTH1) which is related to cellular iron homeostasis. The cluster to the upper right side of the major network covers proteins with major functions mostly related to fatty acid degradation (i.e. Propionyl-CoA carboxylase beta chain, mitochondrial (PCCB), Alpha-aminoadipic semialdehyde dehydrogenase (ALDH7A1), Glutaryl-CoA dehydrogenase, mitochondrial (GCDH)) and amino acid catabolism in mitochondria (i.e. 2-oxoisovalerate dehydrogenase subunit beta, mitochondrial (BCKDHB)) and a redox factor which is used during respiration in electron transport chain (Fumarylacetoacetate hydrolase domain containing 2A (FAHD2A)).

Proteins which are found to be up-regulated in poor quality eggs formed a network made of two major subclusters in total (**Fig 4 Right panel)**. The first subcluster to the top left covers proteins which are mainly involved in mitochondrial structural proteins (i.e. Ubiquinol-cytochrome c reductase binding protein (UQCRB), Cytochrome b-c1 complex subunit Rieske, mitochondrial (UQCRFS1), Cytochrome c1 (CYC1), and ATP synthase F1 subunit beta (ATPF5B)), complex assembly factors (i.e. Complex I assembly factor ACAD9, mitochondrial (ACAD9), NADH:ubiquinone oxidoreductase complex assembly factor 4 (NDUFAF4)), mitochondrial energy generation related proteins (i.e. Fumarate hydratase, mitochondrial (FH), ATP citrate lyase (ACLY), Fructose-bisphosphatase 1 (FBP1), and Glycogen debranching enzyme (AGL)). This cluster is interconnected with two other proteins with the Glycine-tRNA ligase (GARS) which is related to protein translation and the Hsp70-binding protein 1 (HSPBP1) which is related to protein degradation and synthesis inhibition. The second major subcluster to the bottom right covers proteins mainly related to mRNA biogenesis and transcription (i.e. Cleavage stimulation factor subunit 2 (CSTF2), Splicing factor 3b subunit 2 (SF3B2), RNA-binding protein FUS (FUS), Pre-mRNA processing factor 8 (PRPF8), Heterogeneous nuclear ribonucleoprotein A1 (HNRNPA1) and TAR DNA binding protein (TARDBP)). Some other proteins within this cluster are the Mitochondrial import receptor subunit TOM34 (TOMM34) and Kinectin 1 (KTN1) which are involved in mitochondrial biogenesis, the Karyopherin subunit alpha 2 (KPNA2) a nuclear protein import protein, GCN1 activator of EIF2AK4 (GCN1) related to protein degradation and synthesis inhibition and the SET nuclear proto-oncogene (SET) involved in DNA replication and chromatin binding.

Enrichment results for the revealed networks are given in **Table S3**. Aside from being in complete accordance with GO enrichment analyses for biological processes, molecular functions and cellular components results have shown interesting KEGG and Reactome pathway enrichment signatures. Proteins down-regulated in poor quality eggs, on one hand, were enriched in metabolic pathways, RNA degradation, valine, leucine, and isoleucine degradation, fatty acid degradation, necroptosis and glycolysis/gluconeogenesis KEGG pathways and the metabolism Reactome pathway (PPI network enrichment value *p* = 1.70 x 10^-9^). Proteins up-regulated in poor quality eggs, on the other hand, were enriched in Alzheimer’s disease, Parkinson’s disease, thermogenesis, metabolic pathways, oxidative phosphorylation, cardiac muscle contraction, Huntington’s disease, citrate cycle (TCA cycle), spliceosome and non-alcoholic fatty liver disease (NAFLD) KEGG pathways, and the citric acid (TCA) cycle and respiratory electron transport, respiratory electron transport, ATP synthesis by chemiosmotic coupling, and heat production by uncoupling proteins, respiratory electron transport, processing of capped intron containing pre-mRNA, mRNA splicing - major pathway, metabolism, ISG 15 antiviral mechanism, and metabolism of RNA Reactome pathways (PPI network enrichment value *p* = 0.000574).

Taking into account the overall results obtained from the TMT labeling based LC-MS/MS quantification, a total of 21 proteins with significant differential abundance between good and poor quality eggs were chosen as candidate markers of egg quality in this study. The thirteen proteins down-regulated in poor quality eggs were chosen to represent the majority of functional categories with a special emphasis to mitochondrial biogenesis and energy metabolism related proteins. And those up-regulated in poor quality eggs were chosen to mostly represent the mitochondrial biogenesis and energy metabolism functional categories. Fold difference in abundance of candidate proteins between good and poor quality eggs varied between 1.07 and 1.85 for poor quality down-regulated proteins and between 1.07 and 4.67 for poor quality up-regulated proteins. Comparisons in abundance of these proteins between good and poor quality halibut eggs are given in **Fig S2**.

### qPCR

Gene expression levels for the 21 candidate marker proteins with significant difference in protein abundance between good and poor quality eggs are given in **Fig S3**. Four out of these 21 genes (*mt-nd5*, *mt-atp6*, *acly1*, and *dhrs9*) exhibited an increase in gene expression with the same tendency to protein abundance, but these differences were not significantly different. Nine of out these 21 genes (*gcdh*, *ppid*, *gatd3a*, *gfm1*, *cap1*, *phb*, *sod1*, *mecr*, and *vdac*) exhibited a converse tendency to protein abundance and these differences were also not significant. Nevertheless, 8 out of 21 genes (*cyc1*, *fh*, *uqcrb*, *gcn1*, *ghitm*, *uqcrfs1*, *fbp1a*, and *atp5f1a*) exhibit a gene expression pattern with similar increasing tendency to protein abundance and significant differences between good and poor quality eggs (Independent samples t-test, *p* < 0.05 followed by Benjamini Hochberg correction for multiple testing, *p* < 0.05).

### PRM based LC-MS/MS

Differential abundance of 8 (MT-ND5, DHRS9, GATD3A, CAP1, GCN1, FBP1, UQCRFS1, GHITM) out of the 21 candidate marker proteins has been validated via parallel reaction monitoring based LC-MS/MS in this study (**Fig S4**). The number of proteins targeted by this method was limited to the availability of peptides that were suitable for use as reference for this study (See Material and Methods section for details). Results revealed all candidate marker proteins, except GHITM, to exhibit the same tendency of regulation as was detected by TMT-labeling based LC-MS/MS. However, only abundances of five candidate proteins (MT-ND5, DHRS9, GATD3A, FBP1, UQCRFS1) were significantly different between good and poor quality eggs. Results were consistently stable in all representative heavy peptides which varied from 1-3 in number of cases per candidate protein. Respectively, FBP1 and UQCRFS1 are up-regulated while MT-ND5, DHRS9 and GATD3A are down-regulated in poor quality eggs (**Fig S4**). Comparison of protein abundance quantification via TMT and PRM based LC-MS/MS applications and gene expression quantification via qPCR for the eight candidate marker proteins are given in **Fig 5**.

**Fig 5.**
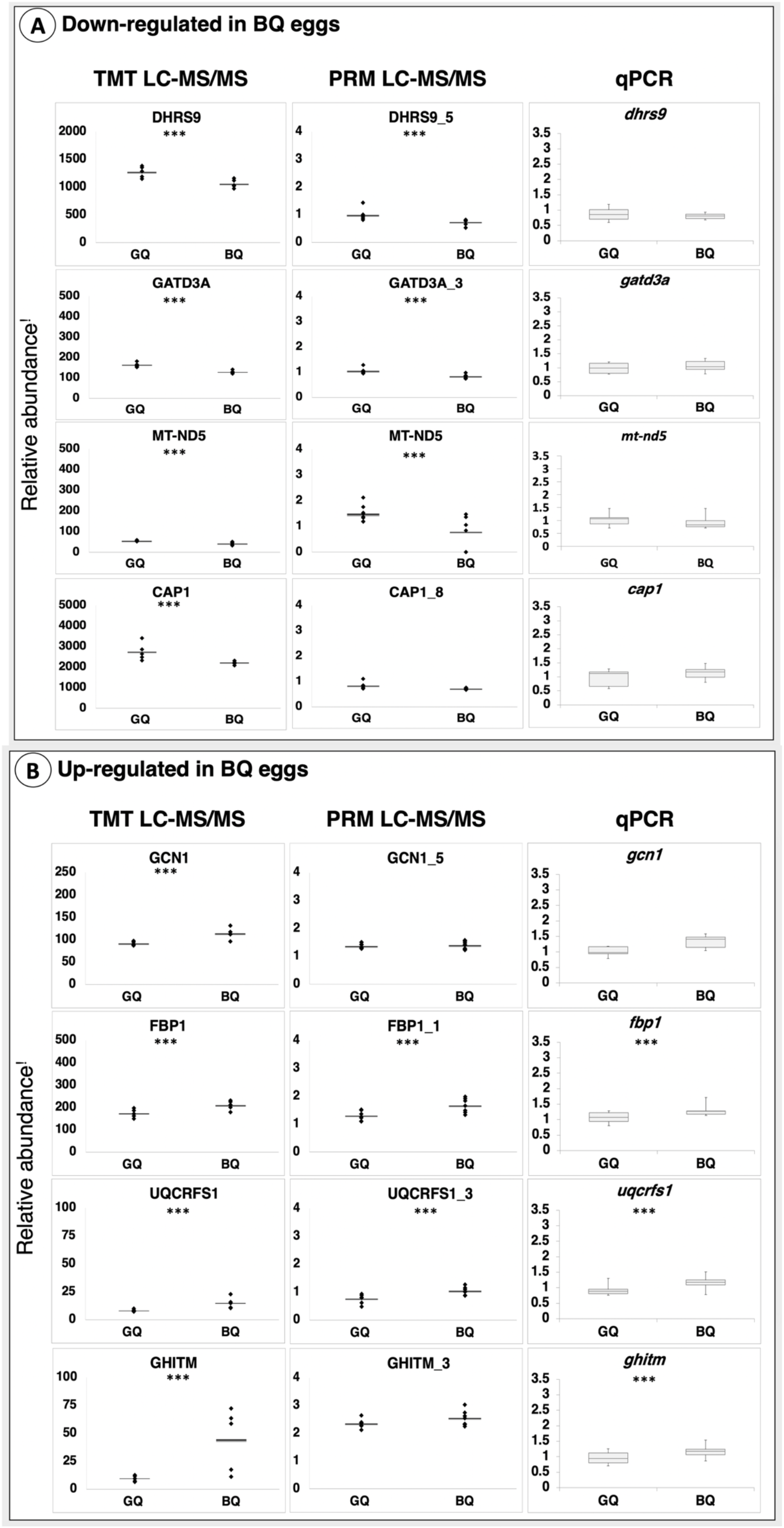
Comparison of marker protein abundances and corresponding gene expressions. **Panel A.** Proteins down-regulated in poor quality eggs **Panel B.** Proteins up-regulated in poor quality eggs. Asterisks indicate significant differences (*p* < 0.05). Relative abundance^!^ represents peak area intensities for protein abundances and gene copy numbers (normalized to transcript copy numbers of halibut *18S*) for gene transcript abundances. GQ: Good quality eggs BQ: poor quality eggs.

### Transmission electron microscopic observations and mtDNA levels

In the guidance of the molecular signatures discovered in this study to be potentially impaired in poor quality eggs, an additional transmission electron microscopy study was conducted with the intention to detect certain morphological differences in mitochondria between good and poor quality eggs. Results, shown in **Fig S5,** revealed the number of vesicles with double membranes which highly resembles intact mitochondria, and the number of intact mitochondria (those with ≥ 5 cristae) to be significantly higher in poor quality eggs (*p* = 0.000724 and *p* = 0.010729, respectively). Accordingly, poor quality eggs seemed to contain about ~1.3 x higher number of vesicles and ^~^1.2 x higher number of intact mitochondria. Poor quality eggs additionally contained significantly higher (1.3 x) cristae number on average in comparison to good quality eggs (*p* = 9.21E-15). Good quality eggs on the other hand, contained larger and well-formed mitochondria with significantly higher mitochondrial area (μm2) and mitochondria circularity (*p* = 1.15E-08 and *p* = 0.016094, respectively). There was no significant difference in total mitochondrial area per cytoplasmic area between good and poor quality eggs (*p* = 0.408). A high variation among females of the same quality group and within eggs from the same batch have been observed. Some eggs from good quality egg batches were observed to contain irregularly shaped empty vesicles (apparently used to be mitochondria) while some others from poor quality egg batches were observed to contain well-formed mitochondria with well-defined cristae. Moreover, some patterns of mitochondrial movements indicating potential to fusion activity were also observed in both good and poor quality eggs. Some examples for these observations are given in **Fig 6 and Fig S6**.

**Fig 6.**
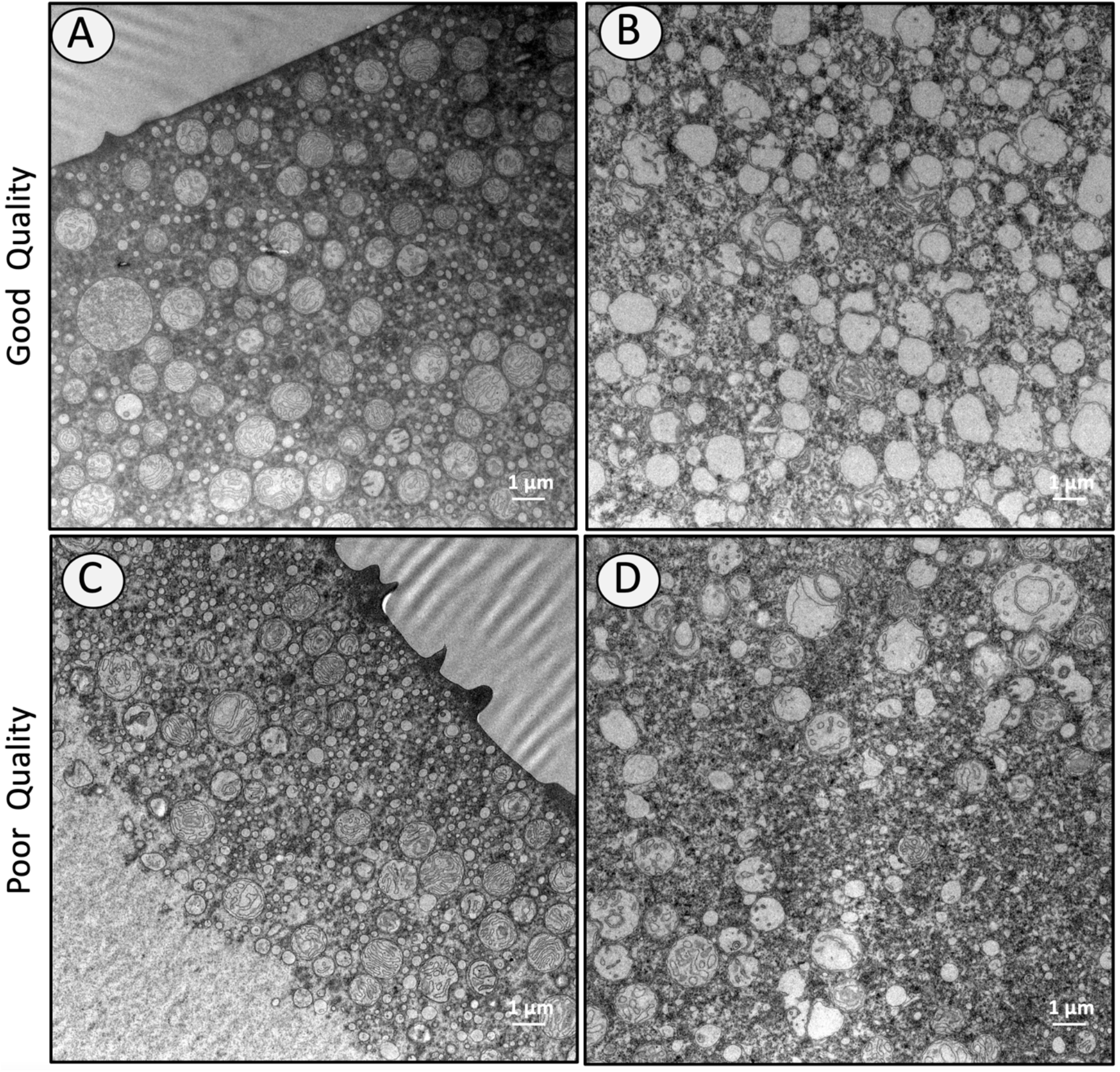
TEM images representing variability of observations between eggs from good and poor quality batches. Despite the standard treatment of biological samples, a high variability has been observed between eggs within the same batch. **Panel A** represents an egg containing a high number of well-formed mitochondria while **Panel B** represents and egg containing a high number of completely deformed mitochondria. Both eggs belonging to the same good quality batch and were kept within the same tube during fixation and postfixation treatments. **Panels C** and **D** on the contrary represents an egg containing a number of better-shaped mitochondria while **Panel B** represents and egg containing a deformed mitochondria. Both eggs belonging to the same poor quality batch and were similarly kept within the same tube during fixation and postfixation treatments. Scalebars indicate 1 μm at 8K magnification.

Significantly higher numbers of smaller and poorly formed mitochondria containing higher number of cristae in poor quality halibut eggs led us to quantify the genomic mitochondrial DNA levels (*mt-nd5* and *mt-atp6*) in good versus poor quality eggs. Results did not reveal any statistically different mtDNA levels in poor quality eggs in comparison to good quality eggs at both 1 hpf and 24 hpf stages (*p* > 0.05) (**Fig 7)**.

**Fig 7.**
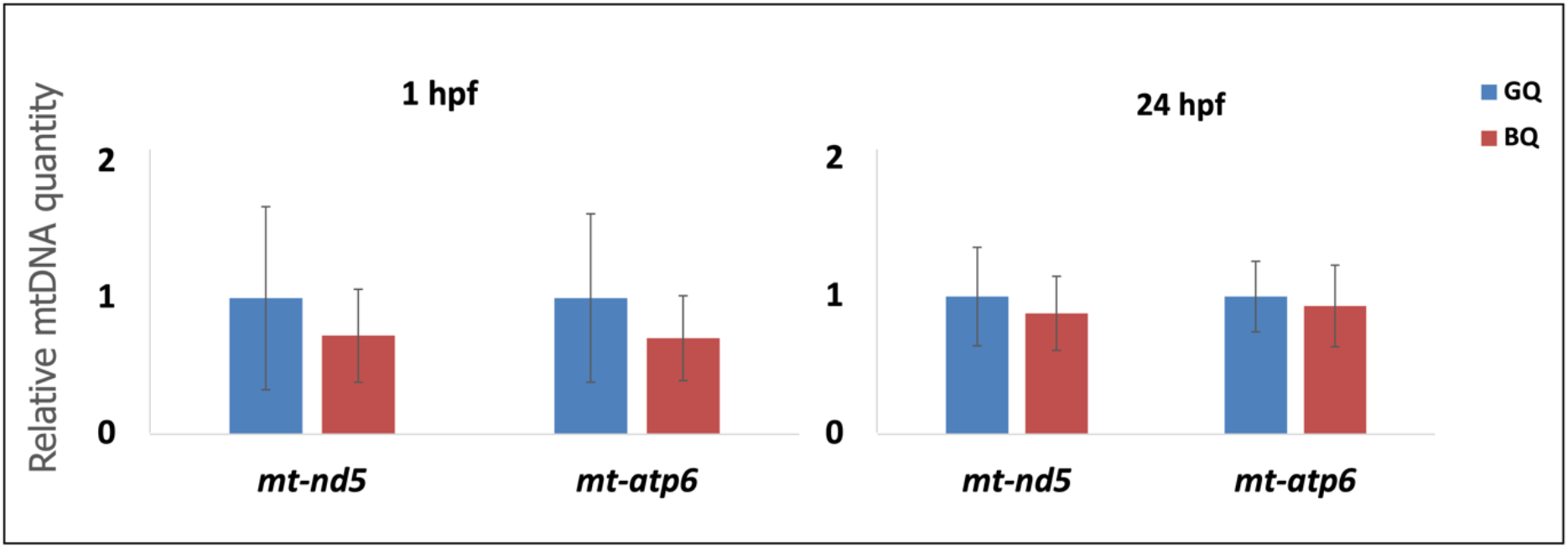
Mitochondrial DNA Quantification. Genomic DNA abundance for *mtnd5* and *mt-atp6* was measured via TaqMan qPCR using standard curve method in 1 hpf and DDCT method with 18S ribosomal RNA as reference gene in 24 hpf halibut eggs. Results indicate no statistically different abundances of mtDNA in poor quality eggs in comparison to good quality eggs (*p* > 0.05) at both stages. GQ: Good quality eggs BQ: poor quality eggs.

## DISCUSSION

### Overview of the biological status of Atlantic halibut eggs

The present study was undertaken to gain insight into the molecular mechanisms involved in egg quality determination in Atlantic halibut. The 1-cell-stage embryo was chosen as the biological material for this study to allow potential comparisons with results obtained from zebrafish in previous studies (22,24) (Yilmaz et al., 2017, Yilmaz et al., 2021). Females of various backgrounds (origin, age, size, experience in reproductive activity) were used as source of biological samples to ensure coverage for multiple factors involved in egg quality determination. An egg quality assessment protocol was established based on embryo survival prior to hatching after correlation assessment among all considered parameters (female fecundity, egg buoyancy, fertilization rate, normal cell division, survival prior hatching) based on experience in hatchery practices. In contrast to other marine species, halibut has shown no clear relation of egg buoyancy to embryo survival. Biological samples which were collected during consecutive reproductive seasons (2019-2021), with slight changes in the set of females, allowed validation of our quality assessment protocol as well as our findings at proteomic and transcriptomic level. The narrow window in survival rate differences between good and poor quality egg batches (14, 5, and 15 %, for 2019, 2020, and 2021, respectively) empowers the significance of our findings. Furthermore, using human database for protein identification, enrichment analyses and protein network analyses fortified the significance of our findings. The overall findings of this study suggest potential disruptions in protein and energy homeostasis mechanisms in poor quality eggs.

### Protein homeostasis

Cellular functions during embryogenesis rely on proteostasis, defined mainly by the appropriate regulation of protein synthesis, protein folding, and protein degradation (25) [Buszczak et al., 2014]. The precise level of protein synthesis that might be ongoing in early stages of embryonic development in fishes is unknown. However, correct protein translation and folding is a crucial step in protein synthesis since accumulation of misfolded and/or unfolded proteins in the ER lumen disturbs its functioning, leading to ER stress which might have severe consequences in developmental competence.

Overall observations from this study suggest somehow blocked or improper translation and protein folding activities in poor quality halibut eggs. Accordingly, the global proteomic profiling results indicate impaired proteostasis in poor quality halibut eggs. Higher frequency distribution of proteins related to protein translation and folding in good quality eggs but of proteins related to transcription and protein degradation and synthesis inhibition in poor quality eggs empowers the signals for this impairment. Additional results with overrepresentation of proteins related to protein folding, RNP complex biogenesis, and RNA catabolic processes in good quality eggs in contrast to overrepresentation of proteins related to RNA splicing and mRNA processing indicate the presence of ER stress conditions and activated UPR mechanisms in poor quality eggs. The absence of a closely interlinked protein homeostasis network in comparison to good quality eggs and down-regulation of proteins related to protein synthesis (PDIA4, PPID, GFM) in poor quality eggs strengthen this hypothesis. These findings strikingly resemble the ones reported for poor quality zebrafish eggs (22) (Yilmaz et al., 2017) and for eggs from females lacking type I and type III vitellogenin genes in their genomes (24) (Yilmaz et al., 2021). Even though a strong connection between the proper function of the multiple Vtg system and egg quality has never been established (26,27) (Pousis et al., 2018, Yilmaz et al., 2018), these common signatures between two evolutionary distinct species express the need for investigations targeting the link in a higher number of fish species in future studies.

Despite the need for more detailed studies on ER stress signaling and UPR, our current findings set a hallmark step taken towards understanding potential impairments of these mechanisms in poor quality fish eggs. Diversely, heat, osmotic and pH stress, maternal nutrition and physiology, ovarian oxidative stress, oxygen and glucose availability and limitations in fatty acid availability are listed among the main factors inducing ER stress and activation of UPR and ER stress signaling in oocytes and embryos of several mammals, including mice, pigs, bovine, rabbit, and human (28) (Latham 2015). The multifariousness in the background of females included in this study fortifies the homogeneity of our findings but it makes it difficult to infer the main causes of the identified impairments. Further research is clearly needed to determine and ascertain the potential causes of these observations.

### Energy homeostasis and mitochondrial biogenesis

Biological activities supporting cell divisions in newly fertilized embryos of egg laying animals are mainly dependent on maternal transcripts, proteins, lipids, and other key molecules loaded into the oocyte prior to final maturation and ovulation. The high amount of energy required to conduct these activities is mainly provided by a normal functioning mitochondria pool, which is produced during oogenesis and peaks during later stages of folliculogenesis (29,30) (St John 2014, Babayev and Seli, 2015). Before embryonic mitochondria take over, the embryo is dependent on the functioning of the existing maternal mitochondria supply to provide the required energy for viability (31,32) [Artuso et al., 2012, Chappel et al., 2013]. Deficiencies in mitochondrial structure and function have been shown to impact egg quality and developmental competence in a vast array of species including human (30,33) [Babayev and Seli, 2015, Ge et al., 2012].

Overrepresentation of mitochondrial biogenesis proteins in poor quality eggs in addition to the specific network mainly formed by proteins related to mitochondrial biogenesis, organization and energy homeostasis were considered as indicators of deteriorations in mitochondrial activities in poor quality eggs in this study. Additional results indicating differential abundance of several mitochondria biogenesis and energy homeostasis related proteins between good and poor quality halibut eggs (MT-ND5, GATD3A, PHB, ACLY, CYC1, FH, UQCRB, GHITM, UQCRFS1, FBP1, and ATP5F1B) were contemplated as proofs to further scrutinize some of these marker candidates at both proteomic and transcriptomic levels. Interestingly enough, six of these candidate marker proteins (UQCRB, CYC1, UQCRFS1, ATP5F1B, FH, FBP1), were found to be haphazardly falling into the network formed by proteins down-regulated in poor quality eggs. Gene expression levels for all these proteins, and more, were possible to quantify successfully while the availability of appropriate target peptides to be used in PRM based LC-MS/MS limited the number of proteins to be investigated at this level. Nevertheless, validation of these findings in sample sets collected at different reproductive seasons for five of these proteins (MT-ND5, GATD3A, GHITM, UQCRFS1, FBP1) has been accomplished successfully via both PRM based LC-MS/MS and qPCR methodologies.

As a result, abundance of several mitochondrial proteins and their corresponding gene expressions revealed significant differences between good and poor quality egg groups. Lower but non-significant protein and transcript abundances of MT-ND5 and MT-ATP6 in contrast to higher and highly significant protein and transcript abundances of CYC1, UQCRB, UQCRFS1 and ATP5F1B in poor quality eggs is intriguing. Nevertheless, all being key components of the inner membrane differential expression of these proteins and their corresponding genes are indicators of structural and functional impairments in mitochondria.

Highly significant overrepresentation of the amino acid degradation, fatty acid degradation, and glycolysis/gluconeogenesis KEGG pathways in good quality eggs, in contrast to overrepresentation of several human neurodegenerative disease pathways and mitochondrial functions related pathways in poor quality eggs may indicate a potential for the presence of more than one problem at the mitochondrial level: lack of substrates for mitochondria to generate energy in addition to structural deficiencies. Again, causes and factors leading to these potential problems are largely unknown and need more detailed studies to be discovered. Prominent KEGG pathways revealed by network enrichment analyses such as Alzheimer’s disease, Parkinson’s disease, Huntington’s disease, oxidative phosphorylation, and citrate cycle in addition to cardiac muscle contraction are highly consistent with the findings from *vtg* lacking zebrafish eggs (24) (Yilmaz et al., 2021). All these pathways seem to be interconnected (34–38) [Chen et al., 2010, Youle et al., 2012, Rugarli and Langer, 2012, Labbadia et al., 2013, Tublin et al., 2019] and previously reported to be linked to perturbations in mitochondrial maintenance, localization, and activity along with aberrant protein folding (37,39) [Williams and Paulson, 2008, Labbadia et al., 2013], all of whom signatures were observed in our study, leading to subsequent impairments in normal development (38) [Tublin et al., 2019]. The cardiac muscle contraction pathway was previously linked to a cardiac and yolk sac edema phenotype observed in zebrafish eggs lacking certain *vtgs* in their genomes (23,24) (Yilmaz et al., 2019, 2021). Further research targeting morphological observations on development of offspring originating from different quality halibut egg batches are needed.

Significant enrichment and differential abundance of proteins related to mitochondrial biogenesis in poor quality eggs led us to investigate certain mitochondrial parameters at transmission electron microscopic level. The two parameters which were considered to represent the abundance of mitochondria were 1) the number of vesicles which resemble mitochondria (vesicles containing double membranes), and 2) the number of intact mitochondria (vesicles containing ≥ 5 solid cristae) per cytoplasmic area. Both parameters were found to be consistently and significantly higher in poor quality eggs. In addition, the number of cristae per mitochondria was also significantly higher in poor quality eggs. These results seem to be puzzling and controversary to the generally accepted concept stating low numbers of mitochondria and mtDNA copies as indicators of low oocyte quality and embryonic developmental competence in several organisms (13,32,40–42) [Chappel 2013, Diez-Juan et al., 2015, Fragouli et al., 2015; reviewed in Kim and Seli, 2019, Ma et al., 2019]. However, they are in fact in accordance with findings from other studies contradicting the utility of mtDNA copy number as marker for embryonic competence in humans (42–46) (Treff et al., 2017, Victor et al., 2017, Klimczak et al., 2018; reviewed in Kim and Seli, 2019, Scott et al., 2020). To test the potential relation between mtDNA levels and mitochondrial abundance we quantified genomic DNA levels of two key mitochondrial genes (*mt-nd5* and *mt-atp6*) in a separate experiment. Results revealed no statistically significant differences in mtDNA abundances between good and poor quality eggs at both 1 hpf and 24 hpf stages. These findings were in accordance with those from a previous study on transcriptome analysis of egg viability in rainbow trout, *Oncorhynchus mykiss* (13) (Ma et al., 2019). In apparent contrast with the non-significant differences in DNA and transcript abundances of *mt-nd5* and *mt-atp6* the significantly higher transcript and protein abundance of some other mitochondrial proteins is intriguing. Significantly higher transcription activities resulting in high numbers of malformed mitochondria despite the similar mtDNA abundance in poor quality eggs might indicate impairments at gene expression and protein synthesis levels. An increasing number of mitochondria has been proposed to be linked to compensatory response of the cell to mitochondrial mutations leading to impaired function and reduction in energy synthesis (47) (Monnot et al. 2013). Smaller and more irregularly shaped mitochondria in poor quality eggs provide supportive evidence of the potential of mitochondrial structural deformities which might be related to ER stress and protein folding deficiencies. The overall findings of the TEM are consistent and complementary to proteomic and transcriptomic findings in this study however, high variability between females of the same quality group and within eggs from the same female necessitate extension of this study with higher number of replicates in the future.

## CONCLUSIONS

This study provides concrete results on signatures of impairments in protein and energy homeostasis related mechanisms in newly fertilized poor quality Atlantic halibut eggs. Such critical impairments and subsequent cellular dysfunctions are marked with solid results from global proteomic profiling, targeted proteomics, transcript and mtDNA abundance measurements and further TEM observations in biological samples of various background collected during three sequential reproductive seasons. The highly variable background of females used as source to different quality egg batches in this study strengthens the legitimacy of the observed molecular signatures. Moreover, high consistency between findings from this and previous zebrafish research might indicate a common stereotypical sequence of interconnected events influencing developmental competence among fishes, human and other mammals. Additional research may be required to validate the use of proteins identified in this study as egg quality markers in fishes and to expediate the details and the potential of these impairments occurring in different species. Nonetheless, this study will pave the way for future research and will help in acceleration of recent advances in the field of embryonic developmental competence of living organisms.

## MATERIAL and METHODS

**Fig S9** and **Fig S10** summarize the process of sample collection and the implemented experimental design, respectively.

### Animal care and biological samples

Egg samples from N = 10, 8, and 6 batches of Atlantic halibut were collected in 2019, 2020 and 2021 reproductive seasons, respectively. Collected samples were from females with various background. The pool included aged and young females (^~^8 to ≥ 17 yrs), small and large females (weight of 25 - 70 kg, and length of 110 - 167 cm), females originated from the wild and F1 generation which were produced in captivity, females which were newly introduced to the system and those with experience in the system (3 and 12 yrs, respectively), and finally females which interchanged in the quality of eggs they release from year to year and those consistently spawning good or poor quality egg batches every year.

A total 22 mature female and male halibut were kept in 7 m diameter ^~^40000 lt capacity circular tanks with natural daylight conditions and sea water at salinity of 34 ppt, taken from 160 m depth. Water temperature ranged from 7.8 to 9 °C from May to December and then decreased to and kept at 6 °C until the end of the spawning season. Fish were hand-fed with an artificial broodstock diet (VITALIS Cal 22 mm, Skretting, Norway) every other day to satiety, except during the spawning season when appetite was low (February-May).

Females were followed closely at the start of the spawning season to determine the first egg release time point and following that were checked every 36 - 42 h for the onset of the following batch release based on morphological changes in the abdominal region. Eggs from spawns between the 3^rd^ and the 5^th^ batch were targeted in this study to ensure the fine tuning of spawning rhythm and stability of egg quality during the season in each female. Following the predicted ovulation of the targeted batch, which occurs at approximately 72 - 92 h after the release of the previous batch, eggs were stripped from mature females and fertilized with sperm collected immediately after. Replicates of 0.5 ml eggs per spawn were snap frozen in liquid nitrogen at 1-cell stage after fertilization and stored at −80 °C until analysis (**Fig S9)**.

Hundred milliliters of fertilized eggs from each spawn were incubated in 250 l incubators and were kept in darkness, at 6 °C until hatching. Daily care involved removal of dead embryos from the bottom of incubators and measurement of their volume for mortality determination. Egg quality assessments were based on embryo survival prior to hatching at 12 dpf (days postfertilization). Egg batches with embryonic survival rates of ≥ 76 were considered to be of good quality and those spawns with ≤ 62 embryonic survival were considered to be of poor quality in 2019. This ratio was ≥ 76 % and ≥ 70 % for good quality egg batches, and ≤ 71 and ≤ 55 for poor quality egg batches in the years 2020, and 2021, respectively. The list of egg batches collected during each year and egg quality assessment parameters and classifications are given in **Table S1**.

### TMT labeling based LC-MS/MS

Egg samples (0.5 ml) from a total of 10 spawns (N = 5 spawns for good quality, N = 5 spawns for poor quality) collected during the 2019 reproductive season were lysed in 1 ml modified RIPA lysis buffer (pH 7.4) containing; 50 mM Tris, 150 mM NaCl, 1 % NP-40, 1 % SDS, 1 % CHAPS, 0.5 % SDC, 1x protease inhibitor cocktail (cOmplete^™^ ULTRA Tablets, Roche). Sample lysis, protein concentration measurement and sample reduction processes were carried out as indicated by (48) Berge et al. (2019) with the following modifications: Samples were sonicated in 6 steps of 30 sec at 40 % amplitude with 30 sec stops between each followed by 30 min incubation on ice and centrifugation at 16200 x g for 30 min at +4 °C. Protein extracts containing 30 μg of total egg proteins were mixed with lysis buffer in 40 μl total volume and reduced with 4 μl of 100 mM DTT for 1 h at RT (at 10 mM of final concentration). Samples were then alkylated with 6 μl of 200 mM Iodoacetamide (IAA) by incubation in dark for 1h at RT. Alkylated samples were then enhanced using Single-Pot Solid-Phase-enhanced Sample Preparation (SP3) according to the protocol by (49) Hughes et al. (2019). Mix of two types of Sera-Mag SpeedBeads 50 mg/ml (GE Helathcare) was prepared at 75 μg/μl bead concentration in 47 μl of water. Four μl of beads mix at a bead/protein ratio of 10:1 (wt/wt) were added onto each alkylated sample along with 126 μl of 100 % EtOH (to 70 % final EtOH concentration). Following 7 min incubation on a thermomixer at 1000 rpm 24 °C samples were washed 3 times in 80 % EtOH. A MagRack system was used to facilitate removal of liquid without disturbing the beads containing proteins of interest.

Tryptic digestion of proteins was carried out using porcine trypsin (Promega, GmbH, Mannheim, Germany). Trypsin solution prepared in 100 mM Ambic and 1 mM CaCl_2_ at a 0.01 μg/μl concentration and 100 μl added onto each sample for a final concentration of ~1.2 μg trypsin per sample containing 30 μg of total protein (trypsin to sample ration 1:25). Samples containing trypsin were then sonicated twice for 30 sec and incubated for 16 h at 1000 rpm 37 °C. Peptides were then recovered by centrifugation at 13000 rpm for 3 min at RT. A second recovery was performed by washing beads with 0.5 M NaCl via pipetting and 2 x ultrasound sonication for 30 sec and centrifugation at 13000 rpm for 3 min at RT. Second recovery of peptide digests was combined with the previous one and peptide concentration was determined on Nanodrop to check for sufficient recovery. Peptide mixtures were desalted and concentrated on reverse-phase Oasis HLB μElution Plate (Waters Corporation, Manchester, UK) as indicated by (50) Yadetie et al., 2014. Lyophilized peptides mixtures were reconstituted in 52 μl of 100 mM Triethyl ammonium bicarbonate (TEAB) buffer and peptide concentrations were determined on Nanodrop to check for sufficient recovery prior isobaric labeling using TMT10plex^™^ Isobaric Label Reagent Set, 1 x 0.8 mg (ThermoFisher Scientific). Twenty-one μl of each label (^~^0.4 mg) were added onto 50 μl of samples containing ~20 μg of peptide digests. After 1 h incubation at RT, 4 μl of 5 % Hydroxylamine (NH2OH) were added and samples were incubated for an additional 15 min at RT to quench the reaction. All ten vials of samples were combined and approximately 100 μg peptide digests from this mix were fractionated using Pierce High pH Reversed-Phase Peptide Fractionation Kit (ThermoFisher Scientific) according to instructions from manufacturer. All fractions were lyophilized and reconstituted in a mix of 0.5 % Formic acid (FA) and 2 % ACN (at ^~^0.5 μg/μl concentration) prior injection to the LC-MS/MS system, an Ultimate 3000 RSLC system (Thermo Scientific, Sunnyvale, California, USA) connected online to a Q-Excative HF mass spectrometer (Thermo Scientific, Bremen, Germany) equipped with EASY-spray nano-electrospray ion source (Thermo Scientific).

Peptides were separated during a biphasic ACN gradient from two nanoflow UPLC pumps (flow rate of 250 nl/min) on a 25 cm analytical column (PepMap RSLC, 25cm x 75 μm i.d. EASY-spray column, packed with 2 μm C18 beads). Solvent A and B were 0.1 % FA (vol/vol) in water and 100 % ACN respectively. The gradient composition was 5 % B during trapping (5min) followed by 5-7 % B over 0.5 min, 7 - 22 % B for the next 59.5 min, 22 - 35 % B over 22 min, and 35 - 80 % B over 5 min. Elution of very hydrophobic peptides and conditioning of the column were performed during 10 min isocratic elution with 80 % B and 15 min isocratic conditioning with 5 % B, respectively. The eluting peptides from the LC-column were ionized in the electrospray and analyzed by the Q-Excative HF. The mass spectrometer was operated in the data-dependent-acquisition mode to automatically switch between full scan MS and MS/MS acquisition. Instrument control was through Q Excative HF Tune 2.9 and Xcalibur 4.1.

MS spectra were acquired in the scan range 375 - 1500 m/z with resolution R = 60 000 at m/z 200, automatic gain control (AGC) target of 3e6 and a maximum injection time (IT) of 50 ms. The 12 most intense eluting peptides above intensity threshold 50 000 counts, and charge states 2 to 6 were sequentially isolated to a target value (AGC) of 1e5 and a maximum IT of 110 ms in the C-trap, and isolation width maintained at 1.6 m/z (offset of 0.3 m/z), before fragmentation in the HCD (Higher-Energy Collision Dissociation) cell. Fragmentation was performed with a normalized collision energy (NCE) of 32 %, and fragments were detected in the Orbitrap at a resolution of 60 000 at m/z 200, with first mass fixed at m/z 110. One MS/MS spectrum of a precursor mass was allowed before dynamic exclusion for 30 sec with “exclude isotopes” on. Lock-mass internal calibration (m/z 445.12003) was used. The spray and ion-source parameters were as follows. Ion spray voltage of 1800 V, no sheath and auxiliary gas flow, and a capillary temperature of 275 °C conditions were additionally set for data acquisition.

### Data Search

Obtained spectra searched against an in-house built proteome database originated from halibut egg transcriptome with additional peptide sequences for mitochondrial proteome and the vitellogenin proteins from this species. Data search was performed using the SequestHT search engine implemented in Proteome Discoverer 2.4 (Thermo Fisher Scientific). Trypsin was selected as protease with a maximum of two missed cleavage sites and cysteine carbamidomethylation and TMT10plex mass tags both at peptide N-terminus and Lysine side chain as fixed modifications. Methionine oxidation was selected as variable modification with a maximum of three such modifications per peptide. The precursor mass tolerance threshold was 10 ppm and the maximum fragment mass error 0.02 Da. A signal-to-noise filter of 1.5 was applied for precursor ions, and only charge states from two to five were used in the search. Filtering out the false positive peptide identifications were performed by means of False Discovery Rate (FDR) on the reversed database, estimated using the Percolator algorithm (http://per-colator.com). Peptide hits were filtered for an FDR of q < 0.01. In addition to the FDR filter, high confident threshold score filters for Sequest HT (cross correlation scores, XCorr) were as follows: 1.9 (z = 2), 2.3 (z = 3), 2.6 (z = 4 or higher). Only proteins/protein groups that were identified by two or more independent peptide hits were accepted as true positive identifications. Proteins that contained similar peptides and could not be differentiated based on MS/MS analysis alone were considered an equivalence class by using the protein grouping algorithm. Only master proteins from each group were considered for the following quantification analysis. Common laboratory contaminants (keratin and albumin proteins) were removed prior to following analysis. The mass spectrometry proteomics data have been deposited to the ProteomeXchange Consortium via the PRIDE (51) (Perez-Riverol et al., 2019) partner repository with the dataset identifier PXD029894 and a project DOI number of 10.6019/PXD029894.

### Data Analysis

Detected proteins were mapped against a common database for all organisms with available correspondent sequences and were identified based on their identities. Protein abundances were quantified based on peak area intensities. Accordingly, differentially abundant proteins were determined based on *p* values resolved from independent samples t-test (*p* < 0.05) followed by Benjamini Hochberg correction for multiple testing (*p* < 0.05) using the SPSS software (IBM SPSS Statistics Version 19.0.0, Armonk, NY). Functional annotation of proteins found to be differentially abundant between good and poor quality eggs was performed using the UNIPROT and KEGG functional annotation tools. These proteins were then classified into thirteen arbitrarily chosen functional categories that would account for ≥ 90 % of the proteins as originally suggested by (22) Yilmaz et al., 2017 with slight modifications. These functional categories are: transcription, translation, protein folding, protein transport, energy metabolism, mitochondrial biogenesis, cell cycle, division, growth and fate, lipid metabolism, metabolism of cofactors and vitamins, protein degradation and synthesis inhibition, oxidoreductase (redox)- and detoxification (detox)-related, and immune response-related. Differentially abundant proteins that could not be attributed to any of these categories and were placed in the category “Other”. For simplicity, proteins were attributed to only one category considered as the ‘best’ fit. Presented results are based on consensus annotations of two independent observers made before any other analyses categorizing the proteins (i.e. observations made ‘blind’). Chi square analysis with significance level of (p ≤ 0.05) was used to detect differences between groups in the distribution of differentially regulated proteins among functional categories.

Gene ontology overrepresentation analyses were conducted using the GESTALT (WEB-based GEne SeT AnaLysis Toolkit) (52) [Liao et al., 2019] available online at for Biological Process, Molecular Function, and Cellular Components, and KEGG Pathway terms using human proteins as reference database. Proteins which were differentially regulated between good and poor quality halibut eggs were additionally subjected to the analysis of protein-protein interaction networks (53) (Szklarczyk et al., 2015) separately using the STRING Network search tool available from the STRING Consortium online at https://string-db.org/cgi/input?sessionId=b1QVfHtqmBW4&input_page_active_form=multiple_identifiers with the data settings Confidence: Medium (0.40), Max Number of Interactions to Show: None/query proteins only. For the GESTALT and STRING analyses, only statistically significant enrichment results (*p* < 0.05) are reported.

### TaqMan based quantitative real time PCR

Gene expression for a total of 21 proteins were tested in good versus poor quality halibut eggs using TaqMan based quantitative real-time PCR (qPCR). Total RNA extraction from frozen N = 19 egg batches, collected from 2019 and 2020 seasons, was performed using TRI Reagent^™^ (Thermo Fisher Scientific). cDNA was synthesized using SuperScript^™^ VILO^™^ cDNA Synthesis Kit (Thermo Fisher Scientific) from 1 μg of DNAse treated (DNase I, Amplification Grade, Thermo Fisher Scientific) total RNA with 260/280 absorbance ratios of 1.9-2.1 (Nanodrop Spectrophotometer, Thermo Fisher Scientific) and RNA integrity values of 9-10 (Bioanalyzer, Agilent Technologies). Gene-specific primers and dual-labelled probes (labelled with 6-carboxyfluorescein and BHQ-1, Black Hole Quencher 1 on 5’ and 3’ terminus, respectively) were designed using Eurofins Genomics qPCR assay design tool available online at https://eurofinsgenomics.eu/en/ecom/tools/qpcr-assay-design/ and Integrated DNA Technologies (IDT) PrimerQuest Tool available online at https://eu.idtdna.com/Primerquest/Home/Index. Designed primers were additionally analyzed for secondary structures using IDT Oligo analyzer tool available online at https://eu.idtdna.com/calc/analyzer and produced by Eurofins Genomics. Sequences of these primers and probes used in this experiment are given in **Table S4**.

Each qPCR was performed in triplicates of 10 μl reactions containing cDNA (diluted at 1:100), 400 nM of each primer, 200 nM of hydrolysis probe, and 1x TaqMan Fast Advanced Master Mix (Applied Biosystems, Thermo Fisher Scientific) according to the manufacturer’s instructions in optical plates on a QuantStudio 5 Real-Time PCR system (ThermoFisher Scientific) equipped with 384-well block. No-template controls for each gene were included for each assay. PCR cycling conditions were as follows: 50°C for 2 mins, 95 °C for 20 s, 40 cycles at 95 °C for 1 s followed by an annealing-extension at 60° C for 20 s. The gene expression abundance within a sample set, relative to Atlantic halibut18S, was calculated using the 2^−ΔΔCt^ mean relative quantification method in this study. Obtained data were subjected to independent samples t-test, *p* < 0.05) followed by Benjamini Hochberg correction for multiple tests, *p* < 0.05 (IBM SPSS Statistics Version 19.0.0, Armonk, NY).

### Parallel reaction monitoring based LC-MS/MS

Eight out of a total 21 proteins (MT-ND5, CAP1, DHRS9, GCN1, GHITM, GATD3A, FBP1, UQCRFS1), which were previously determined as differentially abundant between good and poor quality egg batches using the TMT labeling based LC-MS/MS methodology, were carried out for further assessments as potential candidate biomarkers of egg quality in halibut. A parallel reaction monitoring based LC-MS/MS approach was followed in order to validate the differential abundance of these proteins between good and poor quality eggs originated from spawns collected both in 2019 (N = 4 spawns for good quality, N = 4 spawns for poor quality) and in 2020 (N = 4 spawns for good quality, N = 4 spawns for poor quality). Egg samples were processed in the same manner as mentioned above for TMT labeling method until prior to isobaric labeling step. About 2-3 target peptides were selected for each protein based on the following criteria collected from (54–57) (Lange et al., 2008, Liebler and Zimmerman, 2013, Hoofnagle et al., 2016 and Chiva and Sabido, 2017); uniqueness to the target protein, length of 5-26 aa, ^~^50 % hydrophobicity, no PTMs, no missed cleavages, positioned far downstream from N- or upstream from C-terminal, proper fragmentation (more than 3-4 fragment ions with well-defined peaks), peptide spectral matches (PSMs) (min 3), charges (min 2-3) and clear clustering in peptide abundance between good and poor quality eggs (**Fig S11**). Peptide PRM compatibility and hydrophobicity tests were performed using Peptide Synthesis and Proteotypic Peptide Analyzing Tool available online at ThermoFisher Scientific. List of target proteins and their corresponding target peptides are listed in **Table S5**. Target peptides for each of these proteins were purchased in stable isotope labelled synthetic peptides (SIS) form in crude quality from Thermo Scientific. The C-terminal lysine or arginine in the SIS peptides were replaced by isotope labelled lysine (^13^C_6_, ^15^N_2_) or arginine (^13^C_6_, ^15^N_4_), resulting in a mass difference of 8 Da and 10 Da, respectively, to the corresponding endogenous peptide. The SIS peptides were spiked in equal amounts into the digested protein samples, at approximately the same level as the endogenous peptide, prior to desalting with Oasis HLB μElution Plate (Waters). The PRM data was analyzed using Skyline v1.4 (58) [MacLean et al., 2010] with the most abundant transition for quantification. Independent samples t-test was used to detect significant differences in abundance between good and poor quality eggs (*p* < 0.05).

### Transmission Electron Microscopy

Four to five eggs from each egg batch (N = 6 batches) that was collected during the 2021 reproductive season were prefixed in Karnovsky’s fixative (59) (Karnovsky 1965) containing 5 % glutaraldehyde, 2 % paraformaldehyde, and 0.1 M Sodium cacodylate buffer for 24 h to allow fixation of the chorion to facilitate its mechanical removal. Dechorionated egg samples were placed back into Karnovsky’s fixative and transferred to the TEM facility for the consecutive steps of the sample preparation process. Eggs were postfixed in 1 % osmium tetroxide (EMS # 19134) diluted in 0.1 M sodium cacodylate buffer on ice for 1 hour. Samples were then washed in buffer and dehydrated using a graded ethanol series (30 %, 50 %, 70 %, 96 % and 100 %) before being transferred to a 1:1 solution of 100 % ethanol:propylene oxide in which they were incubated for 15 min. Samples were then incubated in 100 % propylene oxide for 15 min before gradually introducing agar 100 resin (AgarScientific R1031)0. Samples were then incubated in a drop of 100 % resin overnight and then placed in molds with fresh 100 % resin at 60°C for 48h to polymerize. Ultrathin sections of approximately 60 nm were collected from N = 5 different regions of each egg representing good or poor quality batches. Images of ultrathin sections at 8K magnification were used to assess the number of vesicles with double membranes (see **Fig S7** for examples) which highly resembles intact mitochondria and the number of intact mitochondria (those with ≥ 5 cristae) per cytoplasm area. Images at 20K magnification were used to assess the morphological differences such as the mitochondrial area (μm2), total mitochondrial area per cytoplasm area (μm2), mitochondria circularity and cristae number per mitochondria in a total of 1200 μm^2^ area for each egg. Mitochondria circularity is calculated as; 4π(Area)/(Perimeter^2), where 1.0 indicates a perfect circle, while 0.0 indicates an elongated shape. A minimum of 50 counts per egg were collected for the cristae number assessment. Independent samples t-test was used to detect significant differences in mitochondrial counts between good and poor quality eggs (*p* < 0.05) using SPSS (IBM SPSS Statistics Version 19.0.0, Armonk, NY).

### Mitochondrial gene quantification by real-time quantitative PCR

Relative abundance of genomic DNA for *mtnd5* and *mt-atp6* was measured via TaqMan qPCR using standard curve method in 1 hpf and DDCT method in 24 hpf halibut eggs. Serial dilutions of a single good quality sample with known DNA concentration were used as a reference for the standard curve method and the 18S ribosomal RNA was used as a reference for relative quantification using DDCT method. For gDNA extraction from 1 hpf eggs the insoluble materials leftover following homogenization in TRI Reagent during RNA isolation was mixed with 300μl of 100% ethanol, tubes were inverted several times and incubated for 3 mins for genomic DNA isolation. The supernatant was removed after centrifugation at 2000 x g at +4 °C and pellets were resuspended in 1 ml of 0.1 M sodium citrate in 10% ethanol (pH 8.5). Samples were incubated for 30 mins at RT mixing occasionally by gentle inversion. The supernatant was discarded after centrifugation for 5 mins at 2000 x g at +4 °C, pellets were resuspended in 1.5 ml 75 % ethanol and incubated for 20 mins by occasionally mixing by gentle inversion. Following centrifugation for 5 mins at 2000 x g at +4 °C pellets were air dried for 5 mins and resuspended in 100μl of water. gDNA extractions from 24 hpf eggs were performed using QIAamp DNA Mini Kit (Qiagen) following the instructions from the manufacturer. DNA concentrations were quantified using a Nanodrop Spectrophotometer (Thermo Fisher Scientific) and each qPCR reaction was performed in triplicates of 10 μl reactions containing 10 ng gDNA for 1 hpf and 40 ng for 24 hpf eggs, 400 nM of each primer, 200 nM of hydrolysis probe, and 1x TaqMan Fast Advanced Master Mix (Applied Biosystems, Thermo Fisher Scientific) according to the manufacturer’s instructions in optical plates on a QuantStudio 5 Real-Time PCR system (ThermoFisher Scientific) equipped with 384-well block. No-template controls for each gene were included for each assay. PCR cycling conditions were as follows: 50°C for 2 mins, 95 °C for 20 s, 40 cycles at 95 °C for 1 s followed by an annealing-extension at 60° C for 20 s. Assay efficiencies were at 98 %, Slope: −3.368, R^2^: 0.999 and 100 %, Slope: −3.307, R^2^: 0.998 for *mt-nd5* and *mt-atp6*, respectively. Obtained data were subjected to independent samples t-test,*p* < 0.05 (IBM SPSS Statistics Version 19.0.0, Armonk, NY). Sequences for primers and probes used in these assays are given in **Table S4**.

## DECLARATIONS

### Ethics approval and consent to participate

The animal study was reviewed and approved by the Norwegian Animal Research Authority (permit number 22921) and the use of these experimental animals was in accordance with theNorwegian Animal Welfare Act.

### Consent for publication

Not applicable

### Availability of data and materials

The mass spectrometry proteomics data have been deposited to the ProteomeXchange Consortium via the PRIDE partner repository with the dataset identifier PXD029894 and a project DOI number of 10.6019/PXD029894.

### Authors’contribution

OY, BN, AW, FB designed the experiments. AMJ, TH, MM, RMJ, OM, EB, ES, LS performed experiments. TF built and provided reference database for proteomics experiments. OY wrote the manuscript with consultation from all authors. All authors read and approved the final manuscript.

## Acknowledgements

Not applicable

## REFERENCES

1. Tarín JJ, García-Pérez MA, Cano A. Assisted reproductive technology results: Why are live-birth percentages so low?: RESULTS OF ASSISTED REPRODUCTIVE TECHNOLOGY. Mol Reprod Dev. 2014 Jul;81(7):568–83.

2. Keefe D, Kumar M, Kalmbach K. Oocyte competency is the key to embryo potential. Fertil Steril. 2015 Feb;103(2):317–22.

3. Kjorsvik E, Meeren T, Kryvi H, Arnfinnson J, Kvenseth PG. Early development of the digestive tract of cod larvae, Gadus morhua L., during start-feeding and starvation. J Fish Biol. 1991 Jan;38(1):1–15.

4. Bobe J, Labbé C. Egg and sperm quality in fish. Gen Comp Endocrinol. 2010 Feb;165(3):535–48.

5. Migaud H, Bell G, Cabrita E, McAndrew B, Davie A, Bobe J, et al. Gamete quality and broodstock management in temperate fish. Rev Aquac. 2013 May;5:S194–223.

6. Aegerter S, Jalabert B, Bobe J. mRNA stockpile and egg quality in rainbow trout (Oncorhynchus mykiss). Fish Physiol Biochem. 2003;28(1–4):317–8.

7. Bonnet E, Fostier A, Bobe J. Microarray-based analysis of fish egg quality after natural or controlled ovulation. BMC Genomics. 2007 Dec;8(1):55.

8. Mommens M, Fernandes JM, Tollefsen K, Johnston IA, Babiak I. Profiling of the embryonic Atlantic halibut (Hippoglossus hippoglossus L.) transcriptome reveals maternal transcripts as potential markers of embryo quality. BMC Genomics. 2014;15(1):829.

9. Chapman RW, Reading BJ, Sullivan CV. Ovary Transcriptome Profiling via Artificial Intelligence Reveals a Transcriptomic Fingerprint Predicting Egg Quality in Striped Bass, Morone saxatilis. Craft JA, editor. PLoS ONE. 2014 May 12;9(5):e96818.

10. Sullivan CV, Chapman RW, Reading BJ, Anderson PE. Transcriptomics of mRNA and egg quality in farmed fish: Some recent developments and future directions. Gen Comp Endocrinol. 2015 Sep;221:23–30.

11. Żarski D, Nguyen T, Le Cam A, Montfort J, Dutto G, Vidal MO, et al. Transcriptomic Profiling of Egg Quality in Sea Bass (Dicentrarchus labrax) Sheds Light on Genes Involved in Ubiquitination and Translation. Mar Biotechnol. 2017 Feb;19(1):102–15.

12. Cheung CT, Nguyen T, Le Cam A, Patinote A, Journot L, Reynes C, et al. What makes a bad egg? Egg transcriptome reveals dysregulation of translational machinery and novel fertility genes important for fertilization. BMC Genomics. 2019 Dec;20(1):584.

13. Ma H, Martin K, Dixon D, Hernandez AG, Weber GM. Transcriptome analysis of egg viability in rainbow trout, Oncorhynchus mykiss. BMC Genomics. 2019 Dec;20(1):319.

14. Tadros W, Lipshitz HD. The maternal-to-zygotic transition: a play in two acts. Development. 2009 Sep 15;136(18):3033–42.

15. Jukam D, Shariati SAM, Skotheim JM. Zygotic genome activation in vertebrates. 2018;33.

16. Groh KJ, Nesatyy VJ, Segner H, Eggen RIL, Suter MJ-F. Global proteomics analysis of testis and ovary in adult zebrafish (Danio rerio). Fish Physiol Biochem. 2011 Sep;37(3):619–47.

17. Chapovetsky V, Gattegno T, Admon A. Proteomics analysis of the developing fish oocyte. In: Babin PJ, Cerdà J, Lubzens E, editors. The Fish Oocyte. Dordrecht: Springer Netherlands; 2007. p. 99–111. Available from: http://link.springer.com/10.1007/978-1-4020-6235-3_4

18. Rime H, Guitton N, Pineau C, Bonnet E, Bobe J, Jalabert B. Post-ovulatory ageing and egg quality: A proteomic analysis of rainbow trout coelomic fluid. Reprod Biol Endocrinol. 2004;10.

19. Crespel A, Rime H, Fraboulet E, Bobe J, Fauvel C. Egg quality in domesticated and wild seabass (Dicentrarchus labrax): A proteomic analysis.:1.

20. Castets M-D, Schaerlinger B, Silvestre F, Gardeur J-N, Dieu M, Corbier C, et al. Combined analysis of Perca fluviatilis reproductive performance and oocyte proteomic profile. Theriogenology. 2012 Jul;78(2):432–442.e13.

21. Kohn YY, Symonds JE, Kleffmann T, Nakagawa S, Lagisz M, Lokman PM. Proteomic analysis of early-stage embryos: implications for egg quality in hapuku (Polyprion oxygeneios). Fish Physiol Biochem. 2015 Dec;41(6):1403–17.

22. Yilmaz O, Patinote A, Nguyen TV, Com E, Lavigne R, Pineau C, et al. Scrambled eggs: Proteomic portraits and novel biomarkers of egg quality in zebrafish (Danio rerio). Jacobs JM, editor. PLOS ONE. 2017 Nov 16;12(11):e0188084.

23. Yilmaz O, Patinote A, Nguyen T, Com E, Pineau C, Bobe J. Genome editing reveals reproductive and developmental dependencies on specific types of vitellogenin in zebrafish (*Danio rerio*). Mol Reprod Dev. 2019 Sep;86(9): 1168–88.

24. Yilmaz O, Patinote A, Com E, Pineau C, Bobe J. Knock out of specific maternal vitellogenins in zebrafish (Danio rerio) evokes vital changes in egg proteomic profiles that resemble the phenotype of poor quality eggs. BMC Genomics. 2021 Dec;22(1):308.

25. Buszczak M, Signer RAJ, Morrison SJ. Cellular Differences in Protein Synthesis Regulate Tissue Homeostasis. Cell. 2014 Oct;159(2):242–51.

26. Pousis C, Mylonas CC, De Virgilio C, Gadaleta G, Santamaria N, Passantino L, et al. The observed oogenesis impairment in greater amberjack *Seriola dumerili* (Risso, 1810) reared in captivity is not related to an insufficient liver transcription or oocyte uptake of vitellogenin. Aquac Res. 2018 Jan;49(1):243–52.

27. Yilmaz O, Patinote A, Nguyen T, Bobe J. Multiple vitellogenins in zebrafish (Danio rerio): quantitative inventory of genes, transcripts and proteins, and relation to egg quality. Fish Physiol Biochem. 2018 Dec;44(6):1509–25.

28. Latham KE. Endoplasmic Reticulum Stress Signaling in Mammalian Oocytes and Embryos: Life in Balance. In: International Review of Cell and Molecular Biology. Elsevier; 2015 [cited 2021 Dec 18. p. 227–65. Available from: https://linkinghub.elsevier.com/retrieve/pii/S1937644815000064

29. St. John J. The control of mtDNA replication during differentiation and development. Biochim Biophys Acta BBA - Gen Subj. 2014 Apr;1840(4):1345–54.

30. Babayev E, Seli E. Oocyte mitochondrial function and reproduction. Curr Opin Obstet Gynecol. 2015 Jun;27(3):175–81.

31. Artuso L, Romano A, Verri T, Domenichini A, Argenton F, Santorelli FM, et al. Mitochondrial DNA metabolism in early development of zebrafish (Danio rerio). Biochim Biophys Acta BBA - Bioenerg. 2012 Jul;1817(7):1002–11.

32. Chappel S. The Role of Mitochondria from Mature Oocyte to Viable Blastocyst. Obstet Gynecol Int. 2013;2013:1–10.

33. Ge H, Tollner TL, Hu Z, Dai M, Li X, Guan H, et al. The importance of mitochondrial metabolic activity and mitochondrial DNA replication during oocyte maturation in vitro on oocyte quality and subsequent embryo developmental competence. Mol Reprod Dev. 2012 Jun;79(6):392–401.

34. Chen H, Chan DC. Physiological functions of mitochondrial fusion. Ann N Y Acad Sci. 2010;1201(1):21–5.

35. Youle RJ, van der Bliek AM. Mitochondrial Fission, Fusion, and Stress. Science. 2012 Aug 31;337(6098):1062–5.

36. Rugarli EI, Langer T. Mitochondrial quality control: a matter of life and death for neurons. EMBO J. 2012 Mar 21;31(6):1336–49.

37. Labbadia J, Morimoto RI. Huntington’s disease: underlying molecular mechanisms and emerging concepts. Trends Biochem Sci. 2013 Aug;38(8):378–85.

38. Tublin JM, Adelstein JM, del Monte F, Combs CK, Wold LE. Getting to the Heart of Alzheimer Disease. Circ Res. 2019 Jan 4;124(1):142–9.

39. Williams AJ, Paulson HL. Polyglutamine neurodegeneration: protein misfolding revisited. Trends Neurosci. 2008 Oct;31(10):521–8.

40. Diez-Juan A, Rubio C, Marin C, Martinez S, Al-Asmar N, Riboldi M, et al. Mitochondrial DNA content as a viability score in human euploid embryos: less is better. Fertil Steril. 2015 Sep;104(3):534–541.e1.

41. Fragouli E, Spath K, Alfarawati S, Kaper F, Craig A, Michel C-E, et al. Altered Levels of Mitochondrial DNA Are Associated with Female Age, Aneuploidy, and Provide an Independent Measure of Embryonic Implantation Potential. Kim SK, editor. PLOS Genet. 2015 Jun 3;11(6):e1005241.

42. Kim J, Seli E. Mitochondria as a biomarker for IVF outcome. Reproduction. 2019 Jun;157(6):R235–42.

43. Treff NR, Zhan Y, Tao X, Olcha M, Han M, Rajchel J, et al. Levels of trophectoderm mitochondrial DNA do not predict the reproductive potential of sibling embryos. Hum Reprod. 2017 Feb 23;1–9.

44. Victor AR, Brake AJ, Tyndall JC, Griffin DK, Zouves CG, Barnes FL, et al. Accurate quantitation of mitochondrial DNA reveals uniform levels in human blastocysts irrespective of ploidy, age, or implantation potential. Fertil Steril. 2017 Jan;107(1):34–42.e3.

45. Klimczak AM, Pacheco LE, Lewis KE, Massahi N, Richards JP, Kearns WG, et al. Embryonal mitochondrial DNA: relationship to embryo quality and transfer outcomes. J Assist Reprod Genet. 2018 May;35(5):871–7.

46. Scott RT, Sun L, Zhan Y, Marin D, Tao X, Seli E. Mitochondrial DNA content is not predictive of reproductive competence in euploid blastocysts. Reprod Biomed Online. 2020 Aug;41(2):183–90.

47. Monnot S, Samuels DC, Hesters L, Frydman N, Gigarel N, Burlet P, et al. Mutation dependance of the mitochondrial DNA copy number in the first stages of human embryogenesis. Hum Mol Genet. 2013 May 1;22(9):1867–72.

48. Berge T, Eriksson A, Brorson IS, Høgestøl EA, Berg-Hansen P, Døskeland A, et al. Quantitative proteomic analyses of CD4+ and CD8+ T cells reveal differentially expressed proteins in multiple sclerosis patients and healthy controls. Clin Proteomics. 2019 Dec;16(1):19.

49. Hughes CS, Moggridge S, Müller T, Sorensen PH, Morin GB, Krijgsveld J. Single-pot, solid-phase-enhanced sample preparation for proteomics experiments. Nat Protoc. 2019 Jan;14(1):68–85.

50. Yadetie F, Karlsen O, Eide M, Hogstrand C, Goksøyr A. Liver transcriptome analysis of Atlantic cod (Gadus morhua) exposed to PCB 153 indicates effects on cell cycle regulation and lipid metabolism. BMC Genomics. 2014;15(1):481.

51. Perez-Riverol Y, Csordas A, Bai J, Bernal-Llinares M, Hewapathirana S, Kundu DJ, et al. The PRIDE database and related tools and resources in 2019: improving support for quantification data. Nucleic Acids Res. 2019 Jan 8;47(D1):D442–50.

52. Liao Y, Wang J, Jaehnig EJ, Shi Z, Zhang B. WebGestalt 2019: gene set analysis toolkit with revamped UIs and APIs. Nucleic Acids Res. 2019 Jul 2;47(W1):W199–205.

53. Szklarczyk D, Franceschini A, Wyder S, Forslund K, Heller D, Huerta-Cepas J, et al. STRING v10: protein–protein interaction networks, integrated over the tree of life. Nucleic Acids Res. 2015 Jan 28;43(D1):D447–52.

54. Lange V, Picotti P, Domon B, Aebersold R. Selected reaction monitoring for quantitative proteomics: a tutorial. Mol Syst Biol. 2008 Jan;4(1):222.

55. Liebler DC, Zimmerman LJ. Targeted Quantitation of Proteins by Mass Spectrometry. Biochemistry. 2013 Jun 4;52(22):3797–806.

56. Hoofnagle AN, Whiteaker JR, Carr SA, Kuhn E, Liu T, Massoni SA, et al. Recommendations for the Generation, Quantification, Storage, and Handling of Peptides Used for Mass Spectrometry–Based Assays. Clin Chem. 2016 Jan 1;62(1):48–69.

57. Chiva C, Sabidó E. Peptide Selection for Targeted Protein Quantitation. J Proteome Res. 2017 Mar 3;16(3):1376–80.

58. MacLean B, Tomazela DM, Shulman N, Chambers M, Finney GL, Frewen B, et al. Skyline: an open source document editor for creating and analyzing targeted proteomics experiments. Bioinformatics. 2010 Apr 1;26(7):966–8.

59. Karnovsky M J. A formaldehyde-glutaraldehyde fixative of high osmolality for use in electron microscopy. J. Cell Biol. 1965; 27:137–8A.

