## Supplementary information for "Molecular mechanisms involved in Atlantic halibut (*Hippoglossus hippoglossus*) egg quality: impairments at transcription and protein folding levels induce inefficient protein and energy homeostasis during early development"

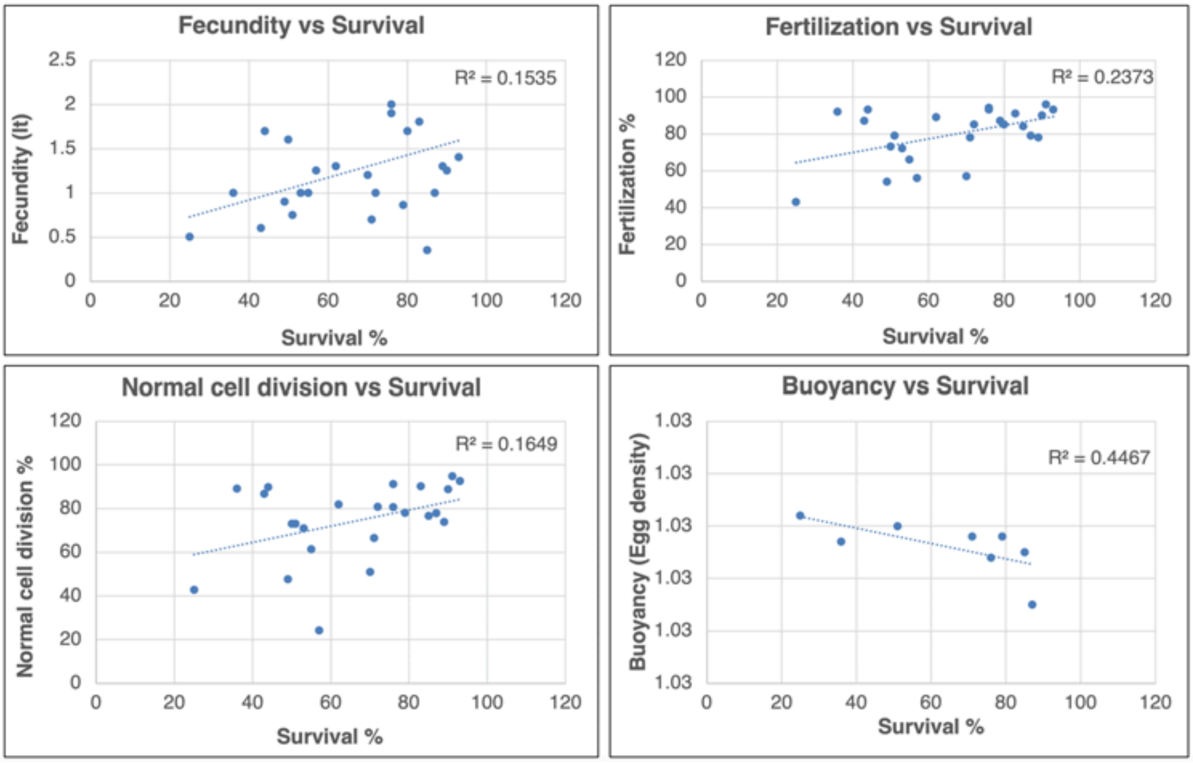

**Fig S1. Correlation of several reproductive parameters with embryonic survival at 12 dpf in halibut. Top left.** Fecundity indicates the volume of eggs (in lt) collected per batch. **Top right.** Fertilization percentage indicates the ratio of embryo with successful visible cell division at 24 hpf. **Bottom left.** Normal cell division percentage represents the number of embryos with symmetrical cell division at 24 hpf. **Bottom right.** Buoyancy indicates egg densities (g/cm$^3$) measured at a buoyancy column at 1 hpf. The $R^2$ values indicating correlation between parameters are given on the top right corner of each panel. Summary of values representing each egg batch used in this study are given in **Table S1**.

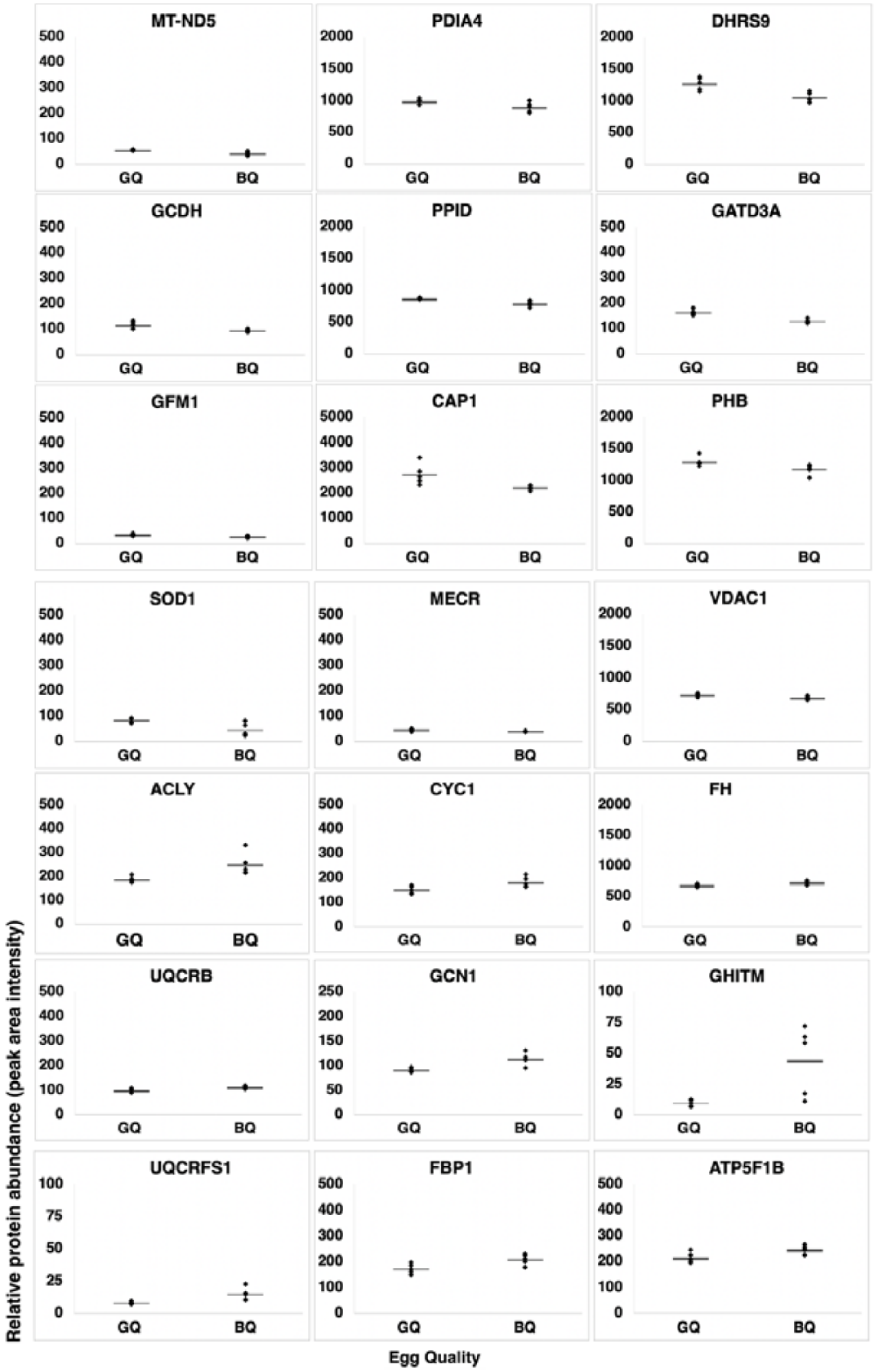

**Fig S2. TMT labeling based LC-MS/MS quantification for candidate marker proteins.** Each panel represents comparisons of protein abundances in peak area intensities between good (GQ) and poor quality (BQ) halibut eggs for 21 candidate marker proteins. All proteins differed significantly in abundance between good and poor quality eggs (Independent t-test $p < 0.05$, followed by Benjamini Hochberg correction for multiple tests $p < 0.05$). See **Table S2** for details of protein names.

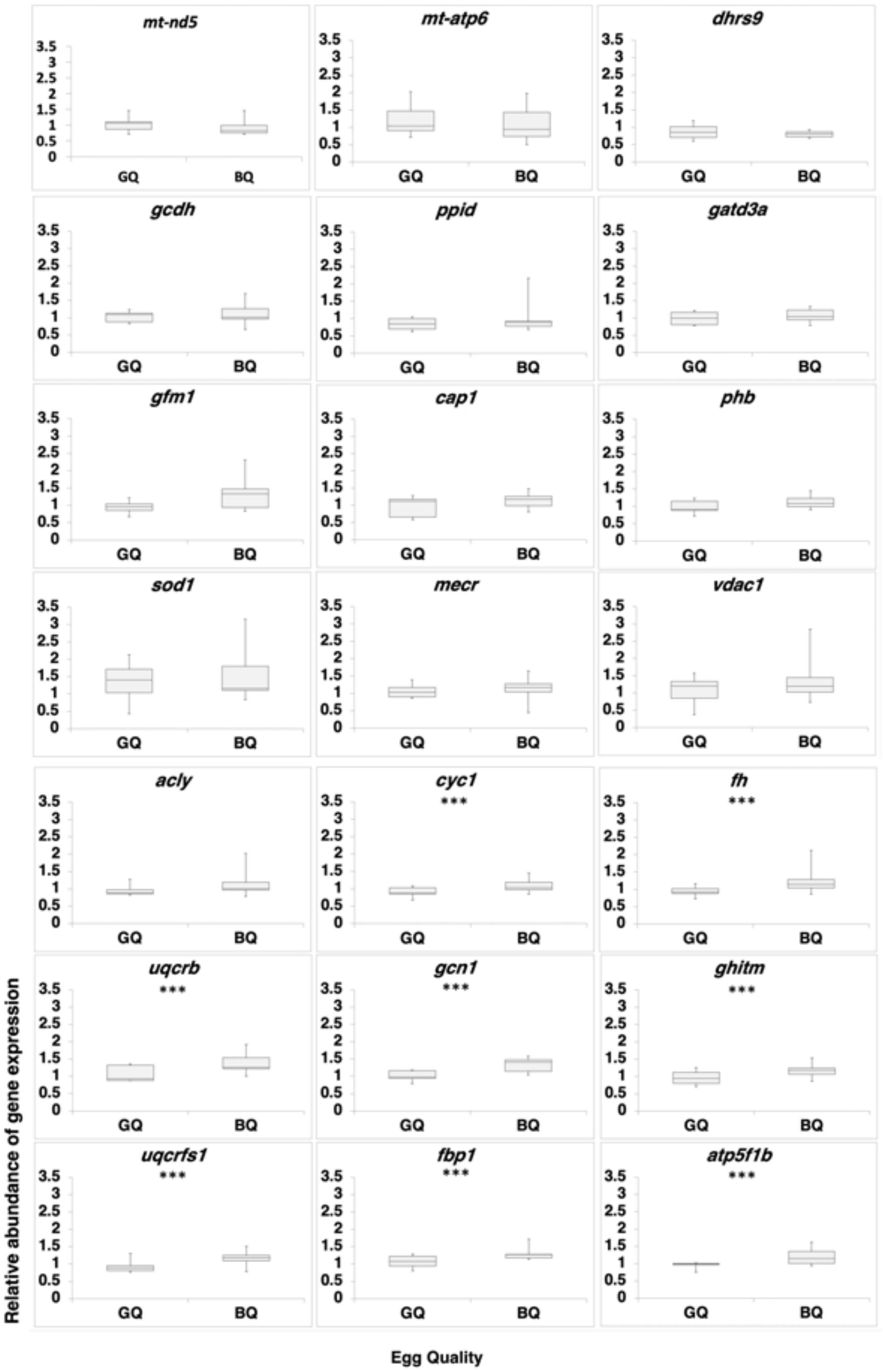

**Fig S3. TaqMan qPCR based relative quantification of gene expression for candidate marker proteins.** Each panel represents comparisons of transcript abundances for 21 differentially abundant proteins between good (GQ) and poor quality (BQ) halibut eggs. TaqMan qPCR-$2^{-\Delta\Delta CT}$ mean relative quantification of gene expression was employed using halibut 18S ribosomal RNA (*18S*) as reference gene. Box plots indicate mean expression ratio (± SEM) with respect to the mean expression of the reference gene. Statistically significant differences are indicated with asterisks (Independent t-test $p < 0.05$ followed by Benjamini Hochberg correction for multiple tests $p < 0.05$).

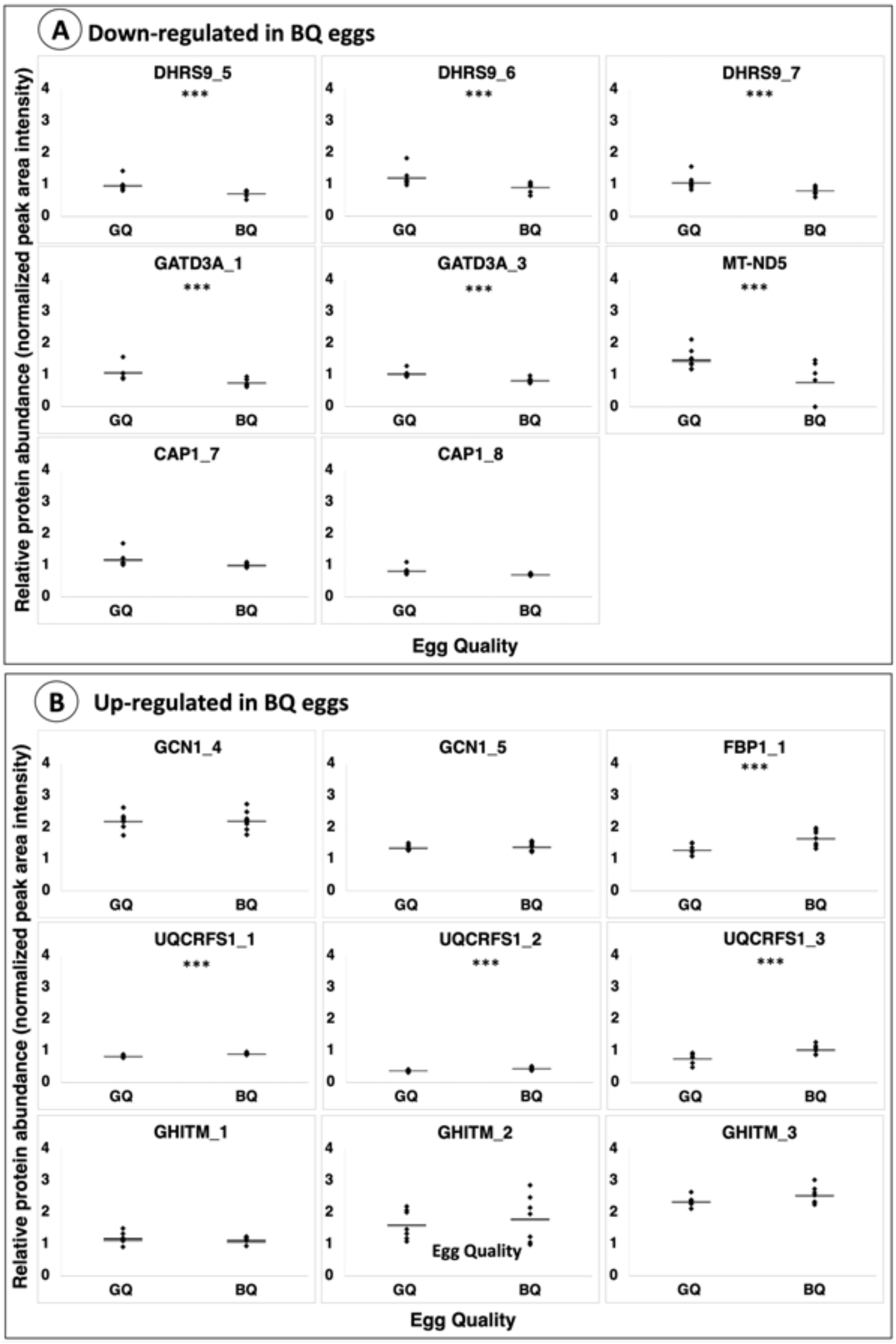

**Fig S4. PRM based LC-MS/MS quantification of protein abundances for candidate marker proteins using**. **Panel A.** Proteins  down-regulated in poor quality (GQ) eggs. **Panel B.** Proteins down-regulate in poor quality (BQ) eggs. Each protein is represented by 1 to 3 reference peptides. The number of candidate marker proteins whom relative abundance has been validated with this method is limted to the avilibility of appropriate reference peptides. Details on selection of reference peptides are given in the **Material and Methods** section and **Fig S9**. Statistically significant differences are indicated with asterisks (Independent t-test $p < 0.05$ followed by Benjamini Hochberg correction for multiple tests $p < 0.05$).

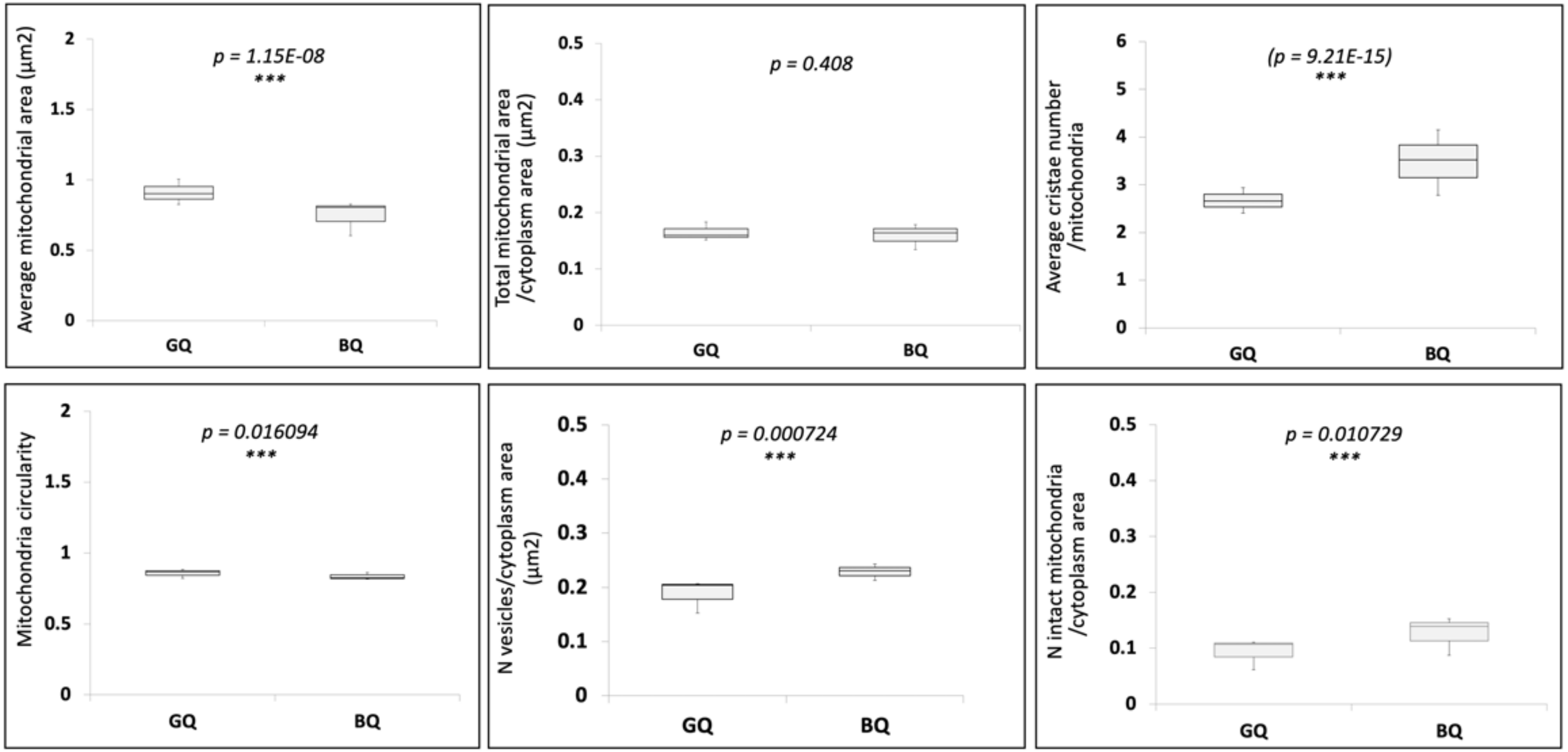

**Fig S5. Results of counts and measurements of some mitochondrial parameters based on TEM imaging.** Each panel represents comparisons between good (GQ) and poor quality (BQ) halibut eggs. Statistically significant differences ($p < 0.05$) are indicated with asterisks and *p* values above each panel.

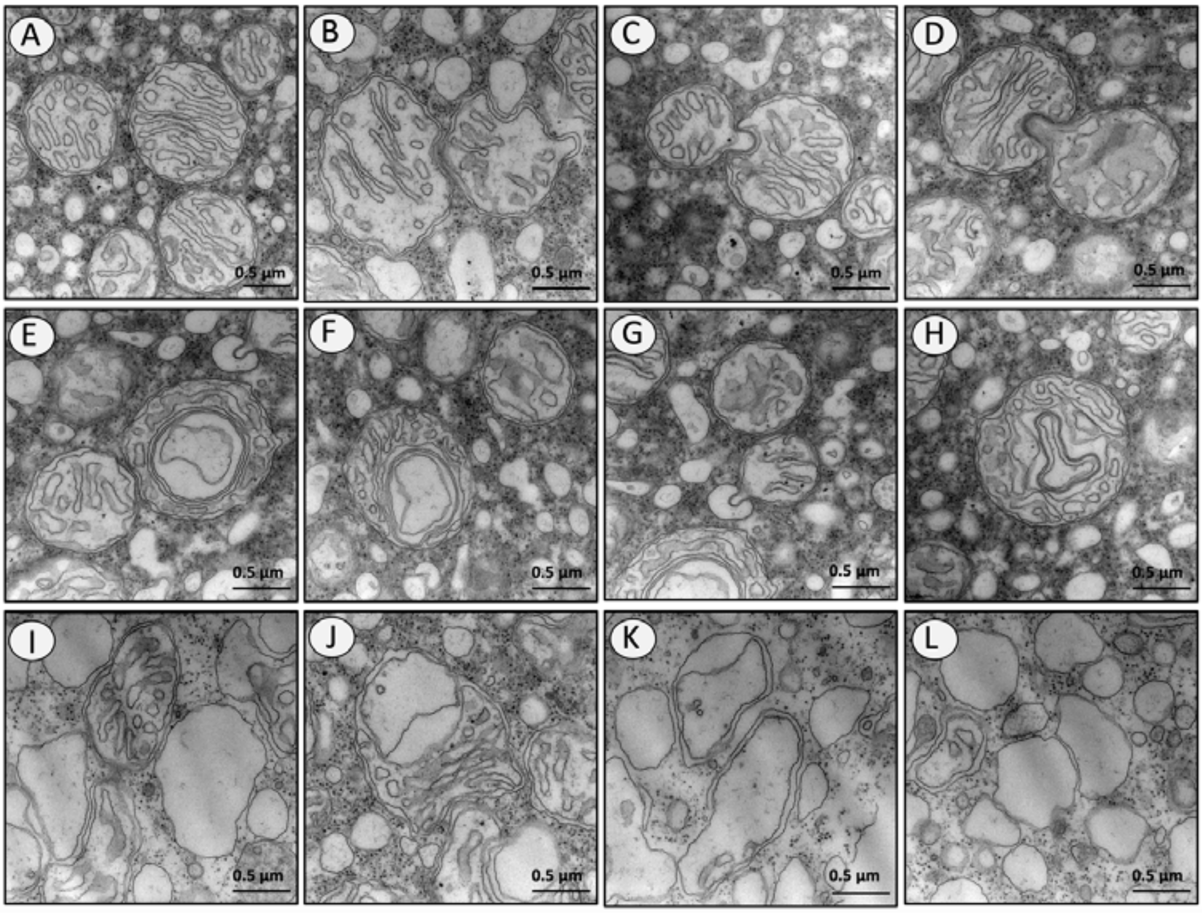

**Fig S6. Some examples of mitochondrial observations in halibut eggs.** Different panels represent different mitochondrial forms and movements that might be related to certain mitochondrial phenomenon observed in this study. **Panel A** shows well-formed examples of mitochondria with different size. **Panels B, C** and **D** show movements of two mitochondria towards fusion. **Panels E, F, G** and **H** exhibit examples of fused mitochondria. **Panels I, J, K** and **L** shows fused or not empty deformed vesicles containing double membranes which assumably used to be mitochondria. Similar mitochondrial events have been observed in both good and poor quality eggs groups. Scalebars indicate 0.5 μm at 50K magnification.

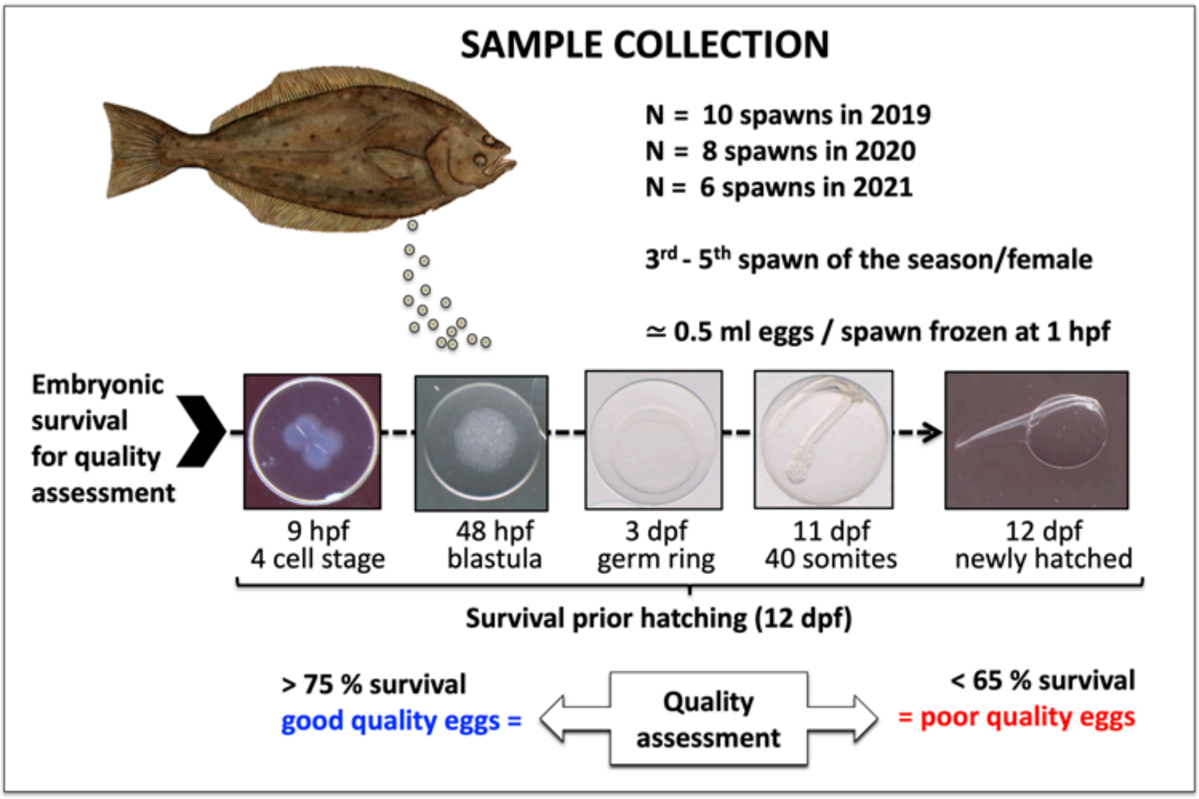

**Fig S7. Biological samples and egg quality assessment procedure used in this study.** A total of 10, 8 and 6 halibut egg batches were collected during the reproductive seasons of 2019, 2020, and 2021, respectively. Between the $3^{rd}$ and the $5^{th}$ batch of each season were targeted to ensure the fine tuning of spawning rhythm and stability of egg quality during the season in each female. Replicates of 0.5 ml eggs per spawn were snap frozen in liquid nitrogen at 1 hpf (at 1-cell stage) and stored at -80 °C until analysis. Hundred milliliters of fertilized eggs from each spawn were incubated until hatching with daily care for mortality determination. Egg quality assessments were based on embryo survival prior hatching at 12 dpf. Egg batches with embryo survival rate of ≥ 76 were considered to be of good quality and those spawns with ≤ 62 embryo survival was considered to be of poor quality in 2019. This ratio was ≥ 76 % and ≥ 70 % for good quality egg batches, and ≤ 71 and ≤ 55 for poor quality egg batches in the years 2020, and 2021, respectively (*See Material and Methods section for details*). The list of egg batches collected during each year and egg quality assessment parameters and classifications are given in **Table S1**. hpf; hours post fertilization, dpf: days postfertilization

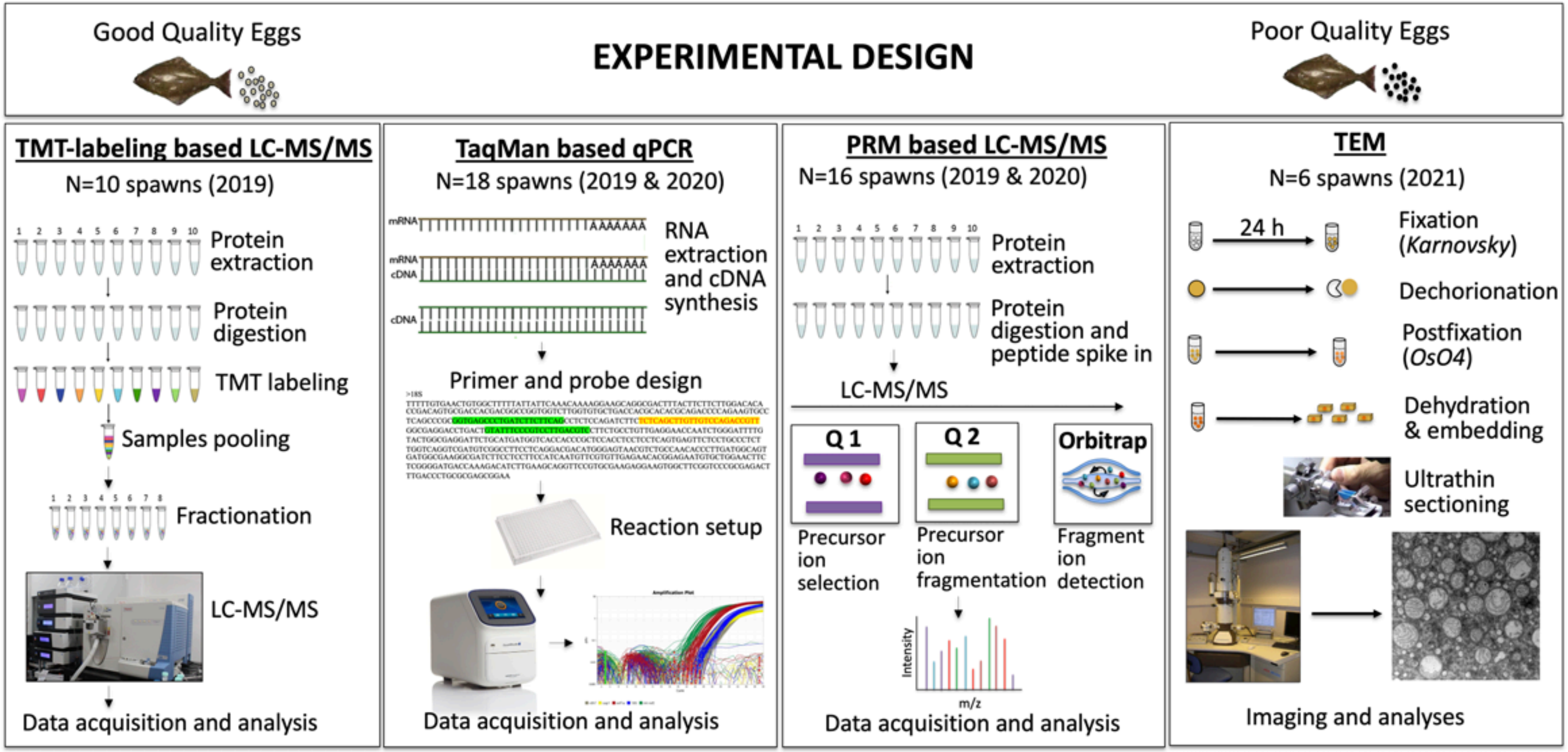

**Fig S8. Experimental design.** Each panel represents the flow of each experimental approach used in this study. The sequence of panels is set in the same order in which each experiment was carried out. Below each panel header the number of biological samples and the year in which they were collected are given.

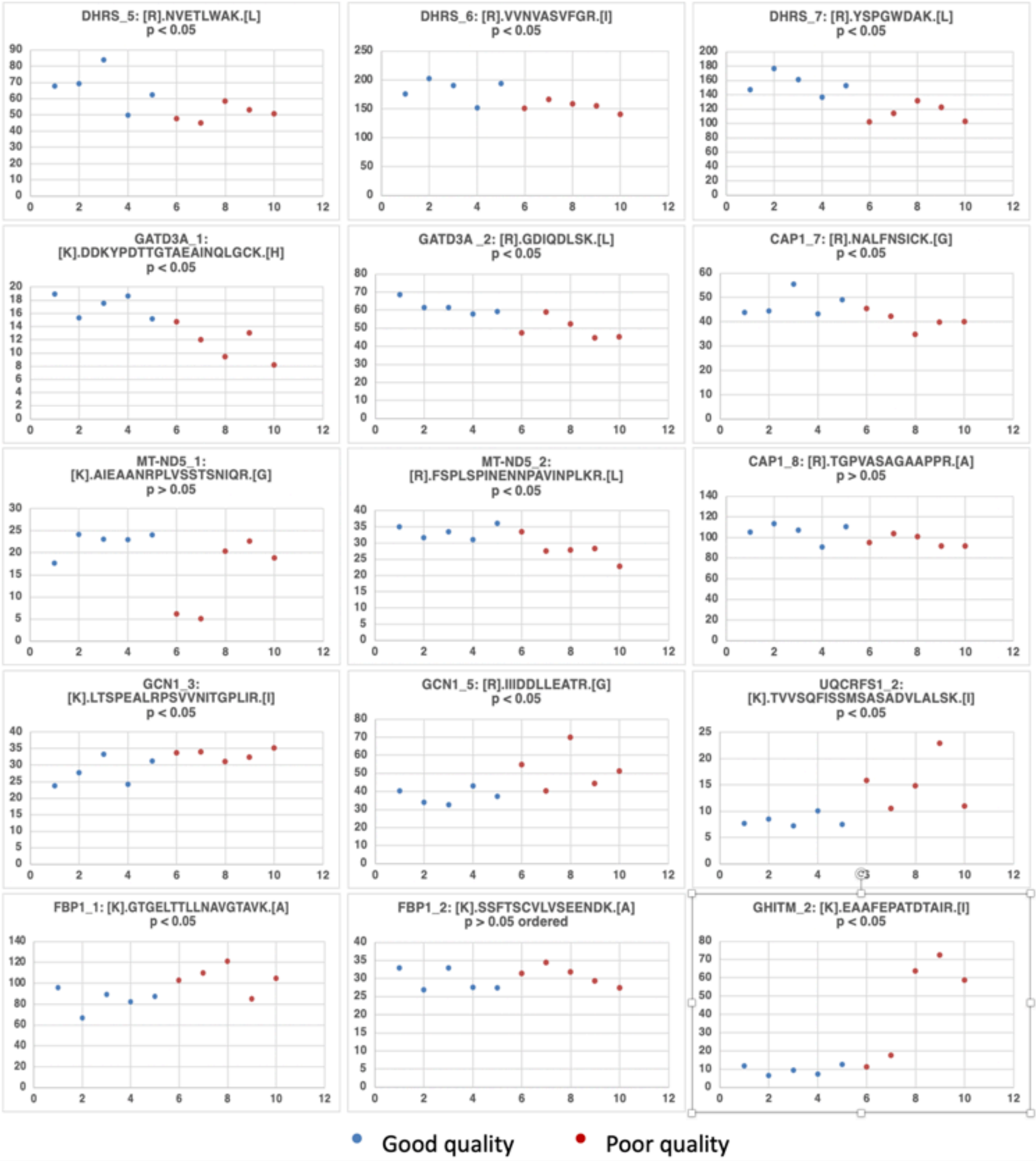

**Fig S9. Quantification of heavy peptides selected for use in PRM based LC-MS/MS.** Each panel contains scatter plots of abundances for each peptide representing differentially abundant proteins. Those results were obtained from the TMT based LC-MS/MS experiment. Blue markers represent values for good quality eggs while red markers indicate values for poor quality eggs. Significance of differences in abundance, the names and the amino acid sequences of each peptide are given above each panel. Clear clustering between good and poor quality eggs along with a *p* value of <0.05 (Independent t-test followed by Benjamini Hochberg correction for multiple testing $p < 0.05$), among other parameters were used for selection of these peptides where possible. See the **Material and Methods** section for details of other criteria used for reference peptide selection in this study.

**Table S1. List of females used in this study.** A list of females from which egg batches were obtained for this study are given. The sampling year, the group and female names, fecundity, fertilization rate, normal cell division rate, egg buoyancy, survival rate at 12 dpf, quality classification, and the application in which the samples were used are given in each column.

| Year | Group | Female name | Fecundity (lt) | Fertilization rate (%) | Normal cell division % | Egg buoyancy (Egg density g/cm3) | Survival rate (%) | Quality | Application |
|---|---|---|---|---|---|---|---|---|---|
| 2019 | Gr3 | Sophia | 1.7 | 93 | 89.96 | | 44 | Poor | TMT, qPCR, PRM |
| 2019 | Gr11 | Sophia | 1.3 | 89 | 81.95 | | 62 | Poor | TMT, qPCR |
| 2019 | Gr12 | Janette | 3.4 | 96 | 94.82 | | 91 | Good | TMT, qPCR, PRM |
| 2019 | Gr15 | PinkFluid BBB0 | 0.9 | 54 | 47.62 | | 49 | Poor | TMT, qPCR, PRM |
| 2019 | Gr17 | Green | 1.3 | 78 | 73.83 | | 89 | Good | TMT, qPCR |
| 2019 | Gr19 | Pink2007 | 1.8 | 91 | 90.23 | | 83 | Good | TMT, qPCR, PRM |
| 2019 | Gr20 | Black | 1.9 | 93 | 80.67 | | 76 | Good | TMT, qPCR, PRM |
| 2019 | Gr21 | Green | 1.4 | 93 | 92.65 | | 93 | Good | TMT, qPCR, PRM |
| 2019 | Gr22 | Red | 1.6 | 73 | 72.94 | | 50 | Poor | TMT, qPCR, PRM |
| 2019 | Gr24 | Rangfrid | 1.25 | 56 | 24.14 | | 57 | Poor | TMT, qPCR, PRM |
| 2020 | Gr2 | Janette | 2 | 94 | 91.18 | 1.0274 ± 0.00002 | 76 | Good | PRM, qPCR |
| 2020 | Gr4 | Sophia | 1 | 92 | 89.09 | 1.0277 ± 0.00001 | 36 | Poor | PRM, qPCR |
| 2020 | Gr8 | PInk2007 | 0.86 | 87 | 77.99 | 1.0278 ± 0.00002 | 79 | Good | PRM, qPCR |
| 2020 | Gr9 | Black | 0.7 | 78 | 66.67 | 1.0278 ± 0.00001 | 71 | Poor | PRM, qPCR |
| 2020 | Gr10 | Ny-liten | 0.5 | 43 | 42.79 | 1.0282 ± 0.00002 | 25 | Poor | PRM, qPCR |
| 2020 | Gr11 | Red | 0.75 | 79 | 73.08 | 1.0280 ± 0.00002 | 51 | Poor | PRM, qPCR |
| 2020 | Gr13 | Yellow | 1 | 79 | 77.91 | 1.0265 ± 0.00002 | 87 | Good | PRM, qPCR |
| 2020 | Gr14 | Green | 0.35 | 84 | 76.72 | 1.0275 ± 0.00001 | 85 | Good | PRM, qPCR |
| 2021 | Gr8 | Pink2007 | 1 | 85 | 80.79 | | 72 | Good | TEM |
| 2021 | Gr9 | Black | 1.2 | 57 | 51.02 | | 70 | Good | TEM |
| 2021 | Gr10 | Red | 1 | 66 | 61.29 | | 55 | Poor | TEM |
| 2021 | Gr11 | Sissel | 0.6 | 87 | 86.70 | | 43 | Poor | TEM |
| 2021 | Gr12 | Yellow | 1 | 72 | 70.95 | | 53 | Poor | TEM |
| 2021 | Gr14 | Ozlem | 1.25 | 90 | 88.78 | | 90 | Good | TEM |

**Table S2. Differentially abundant proteins between good and poor quality halibut eggs based on peak are intensities obtained from the TMT based LC-MS/MS.** List of the 115 proteins which were differentially regulated between good (GQ) and poor quality (BQ) eggs and whose distribution among various functional categories is illustrated in Fig 2. These include proteins up-regulated in poor quality eggs (BQ-upregulated, N = 51) and proteins down-regulated in poor quality eggs (BQ-downregulated, N = 64). For each protein, the NCBI gene IDs, NCBI accession numbers, associated protein names from human database, protein full names, functional categories (Fig 2), significance of differences in abundance (Independent t- test $p < 0.05$, followed by Benjamini Hochberg correction for multiple tests $p < 0.05$), relative abundance ratios (GQ/BQ and BQ/GQ, respectively), and regulation tendencies (BQ-upregulated or BQ-downregulated) are given in each column. Colour shading corresponds to that used to designate functional categories in **Fig 2**.

| | NCBI Gene ID | NCBI Accession Number | Associated Gene Name | Protein Full Name | Functional Category | Significance | GQ/BQ | BQ/GQ | Regulation |
|---|---|---|---|---|---|---|---|---|---|
| 1 | 117752991 | XP_034426650.1 | GHITM | Growth hormone inducible transmembrane protein | Mitochondrial biogenesis | A( .024) | 0.21 | 4.67 | BQ-upreg |
| 2 | 117763568 | XP_034444681.1 | UQCRFS1 | Cytochrome b-c1 complex subunit Rieske, mitochondrial | Mitochondrial biogenesis | A( .018) | 0.55 | 1.83 | BQ-upreg |
| 3 | 117760082 | XP_034438719.1 | NEK1 | NIMA related kinase 1 | Cell cycle, division, growth and fate | A( .015) | 0.55 | 1.80 | BQ-upreg |
| 4 | 117758087 | XP_034435339.1 | NDUFAF4 | NADH:ubiquinone oxidoreductase complex assembly factor 4 | Mitochondrial biogenesis | A( .005) | 0.61 | 1.64 | BQ-upreg |
| 5 | 117762216 | XP_034442625.1 | CAND1 | Cullin associated and neddylation dissociated 1 | Protein degradation and synthesis inhibition | A( .031) | 0.67 | 1.50 | BQ-upreg |
| 6 | | | IGHV4-59 | Immunoglobulin heavy variable 4-59 | Immune response related | A( .016) | 0.68 | 1.47 | BQ-upreg |
| 7 | 117777424 | XP_034468099.1 | RRAGC | Ras related GTP binding C | Protein degradation and synthesis inhibition | A( .007) | 0.75 | 1.34 | BQ-upreg |
| 8 | 117752512 | XP_034425755.1 | ACLY | ATP-citrate synthase isoform X2 | Energy metabolism | A( .023) | 0.75 | 1.34 | BQ-upreg |
| 9 | 117754080 | XP_034428638.1 | PRMT6 | Protein arginine methyltransferase 6 | Cell cycle, division, growth and fate | A( .041) | 0.75 | 1.33 | BQ-upreg |
| 10 | 117770867 | XP_034456538.1 | DENND3 | DENN domain containing 3 | Protein degradation and synthesis inhibition | A( .019) | 0.79 | 1.26 | BQ-upreg |
| 11 | 117762822 | XP_034443349.1 | PCSK5 | Proprotein convertase subtilisin/kexin type 5 | Protein folding | A( .026) | 0.80 | 1.26 | BQ-upreg |
| 12 | 117755772 | XP_034431729.1 | KTN1 | Kinectin 1 | Mitochondrial biogenesis | A( .005) | 0.80 | 1.24 | BQ-upreg |
| 13 | 117767628 | XP_034451313.1 | SET | SET nuclear proto-oncogene | Cell cycle, division, growth and fate | A( .004) | 0.81 | 1.24 | BQ-upreg |
| 14 | 117767474 | XP_034451022.1 | GCN1 | GCN1 activator of EIF2AK4 | Protein degradation and synthesis inhibition | A( .006) | 0.81 | 1.24 | BQ-upreg |
| 15 | 117777807 | XP_034468662.1 | AGL | Amylo-alpha-1, 6-glucosidase, 4-alpha-glucanotransferase a | Energy metabolism | A( .027) | 0.81 | 1.24 | BQ-upreg |
| 16 | 117775126 | XP_034463892.1 | CYC1 | Cytochrome c1 | Mitochondrial biogenesis | A( .026) | 0.82 | 1.21 | BQ-upreg |
| 17 | 117759285 | XP_034437238.1 | NAE1 | NEDD8 activating enzyme E1 subunit 1 | Cell cycle, division, growth and fate | A( .032) | 0.83 | 1.21 | BQ-upreg |
| 18 | 117768103 | XP_034452092.1 | FBP1 | Fructose-1,6-bisphosphatase 1b | Energy metabolism | A( .021) | 0.83 | 1.21 | BQ-upreg |
| 19 | 117763675 | XP_034444914.1 | SCAMP2 | Secretory carrier-associated membrane protein 2 isoform X1 | Protein transport | A( .021) | 0.83 | 1.20 | BQ-upreg |
| 20 | 117761432 | XP_034441219.1 | PDZRN3 | E3 ubiquitin-protein ligase PDZRN3-B | Protein degradation and synthesis inhibition | A( .000) | 0.84 | 1.19 | BQ-upreg |
| 21 | 117769462 | XP_034454252.1 | QDPR | Dihydropteridine reductase | Redox/detox related | A( .028) | 0.84 | 1.19 | BQ-upreg |
| 22 | 117765983 | XP_034448541.1 | FUS | RNA-binding protein FUS | Transcription | A( .019) | 0.85 | 1.18 | BQ-upreg |
| 23 | 117754984 | XP_034430255.1 | TFDP1 | Transcription factor Dp-1 | Cell cycle, division, growth and fate | A( .042) | 0.85 | 1.17 | BQ-upreg |
| 24 | 117759545 | XP_034437610.1 | AP1G1 | Adaptor related protein complex 1 subunit gamma 1 | Protein transport | A( .014) | 0.85 | 1.17 | BQ-upreg |
| 25 | 117761614 | XP_034441558.1 | TARDBP | TAR DNA binding protein | Transcription | A( .026) | 0.86 | 1.16 | BQ-upreg |
| 26 | 117754912 | XP_034430157.1 | ATP5F1B | ATP synthase subunit beta, mitochondrial | Mitochondrial biogenesis | A( .021) | 0.87 | 1.15 | BQ-upreg |
| 27 | 117771789 | XP_034458447.1 | FUBP3 | Far upstream element binding protein 3 | Transcription | A( .023) | 0.87 | 1.15 | BQ-upreg |
| 28 | 117752048 | XP_034425068.1 | ZP4 | Zona pellucida glycoprotein 4 | Others | A( .044) | 0.88 | 1.14 | BQ-upreg |
| 29 | 117752579 | XP_034425926.1 | PPM1B | Protein phosphatase, Mg2+/Mn2+ dependent 1B | Cell cycle, division, growth and fate | A( .021) | 0.88 | 1.14 | BQ-upreg |
| 30 | 117779089 | XP_034470836.1 | SF3B2 | Splicing factor 3b subunit 2 | Transcription | A( .017) | 0.88 | 1.14 | BQ-upreg |
| 31 | 117770151 | XP_034455123.1 | UQCRB | Cytochrome b-c1 complex subunit 7 | Mitochondrial biogenesis | A( .020) | 0.88 | 1.14 | BQ-upreg |
| 32 | 117773394 | XP_034461171.1 | PRPF8 | Pre-mRNA processing factor 8 | Transcription | A( .018) | 0.89 | 1.13 | BQ-upreg |
| 33 | 117766753 | XP_034449984.1 | HSPBP1 | Hsp70-binding protein 1 | Protein degradation and synthesis inhibition | A( .045) | 0.89 | 1.13 | BQ-upreg |
| 34 | 117763422 | XP_034444410.1 | LCAT | Phosphatidylcholine-sterol acyltransferase | Lipid metabolism | A( .026) | 0.89 | 1.12 | BQ-upreg |
| 35 | 117761636 | XP_034441595.1 | TOMM34 | Mitochondrial import receptor subunit TOM34 | Mitochondrial biogenesis | A( .011) | 0.89 | 1.12 | BQ-upreg |
| 36 | 117769197 | XP_034453826.1 | UBXN1 | UBX domain-containing protein 1 | Protein degradation and synthesis inhibition | A( .023) | 0.90 | 1.11 | BQ-upreg |
| 37 | 117764325 | XP_034445894.1 | HNRNPA1 | Heterogeneous nuclear ribonucleoprotein A1 | Transcription | A( .043) | 0.90 | 1.11 | BQ-upreg |
| 38 | 117766247 | XP_034449002.1 | KPNA2 | Karyopherin subunit alpha 2 | Protein transport | A( .031) | 0.91 | 1.10 | BQ-upreg |
| 39 | 117771710 | XP_034458294.1 | NUP188 | Nucleoporin 188 | Cell cycle, division, growth and fate | A( .015) | 0.92 | 1.09 | BQ-upreg |
| 40 | 117769321 | XP_034454063.1 | ATL3 | Atlastin-3 | Protein transport | A( .031) | 0.92 | 1.08 | BQ-upreg |
| 41 | 117768844 | XP_034453131.1 | CSTF2 | Cleavage stimulation factor subunit 2 | Transcription | A( .023) | 0.92 | 1.08 | BQ-upreg |
| 42 | 117757984 | XP_034435135.1 | PREP | Prolyl endopeptidase | Protein degradation and synthesis inhibition | A( .001) | 0.93 | 1.08 | BQ-upreg |
| 43 | 117762157 | XP_034442557.1 | PDCD4 | Programmed cell death protein 4 | Protein degradation and synthesis inhibition | A( .010) | 0.93 | 1.08 | BQ-upreg |
| 44 | 117770435 | XP_034455792.1 | ASAH1 | N-acylsphingosine amidohydrolase 1 | Lipid metabolism | A( .035) | 0.93 | 1.08 | BQ-upreg |
| 45 | 117760820 | XP_034440059.1 | FH | Fumarate hydratase, mitochondrial | Energy metabolism | A( .030) | 0.93 | 1.07 | BQ-upreg |
| 46 | 117761735 | XP_034441806.1 | ACAD9 | Complex I assembly factor ACAD9, mitochondrial | Mitochondrial biogenesis | A( .049) | 0.93 | 1.07 | BQ-upreg |
| 47 | 117771002 | XP_034456885.1 | PITHD1 | PITH domain containing 1 | Transcription | A( .014) | 0.94 | 1.07 | BQ-upreg |
| 48 | 117766721 | XP_034449918.1 | POLD1 | DNA polymerase delta catalytic subunit | Cell cycle, division, growth and fate | A( .047) | 0.94 | 1.06 | BQ-upreg |
| 49 | 117754234 | XP_034428951.1 | GARS | Glycine--tRNA ligase | Translation | A( .041) | 0.95 | 1.05 | BQ-upreg |
| 50 | 117766068 | XP_034448668.1 | TUBB | Tubulin beta chain | Cell cycle, division, growth and fate | A( .030) | 0.96 | 1.04 | BQ-upreg |
| 51 | 117765772 | XP_034448148.1 | CNP | 2',3'-cyclic-nucleotide 3'-phosphodiesterase | Cell cycle, division, growth and fate | A( .008) | 0.97 | 1.03 | BQ-upreg |
| 1 | 117774973 | XP_034463638.1 | GEMIN5 | Gem-associated protein 5 | Translation | B( .029) | 1.04 | 0.97 | BQ-downreg |
| 2 | 117754575 | XP_034429430.1 | DUT | Deoxyuridine 5'-triphosphate nucleotidohydrolase, mitochondrial | Cell cycle, division, growth and fate | B( .015) | 1.05 | 0.95 | BQ-downreg |
| 3 | 117761570 | XP_034441479.1 | RPN2 | Dolichyl-diphosphooligosaccharide--protein glycosyltransferase subunit 2 | Protein folding | B( .034) | 1.06 | 0.95 | BQ-downreg |
| 4 | 117760170 | XP_034438870.1 | THOP1 | Thimet oligopeptidase 1 | Protein degradation and synthesis inhibition | B( .046) | 1.07 | 0.94 | BQ-downreg |
| 5 | 117774375 | XP_034462663.1 | SMPD1 | Sphingomyelin phosphodiesterase 1 | Lipid metabolism | B( .037) | 1.07 | 0.94 | BQ-downreg |
| 6 | 117760379 | XP_034439250.1 | ALDH7A1 | Alpha-aminoadipic semialdehyde dehydrogenase | Energy metabolism | B( .039) | 1.07 | 0.94 | BQ-downreg |
| 7 | 117774288 | XP_034462485.1 | VDAC1 | Voltage dependent anion channel 1 | Cell cycle, division, growth and fate | B( .018) | 1.07 | 0.93 | BQ-downreg |
| 8 | 117768013 | XP_034451932.1 | FAHD2A | Fumarylacetoacetate hydrolase domain containing 2A | Energy metabolism | B( .019) | 1.07 | 0.93 | BQ-downreg |
| 9 | 117768384 | XP_034452554.1 | ACTB | Actin, cytoplasmic | Cell cycle, division, growth and fate | B( .035) | 1.08 | 0.93 | BQ-downreg |
| 10 | 117761895 | XP_034442117.1 | METTL7A | Methyltransferase like 7A | Protein folding | B( .012) | 1.08 | 0.92 | BQ-downreg |
| 11 | 117761802 | XP_034441952.1 | AHCYL1 | Adenosylhomocysteinase like 1 | Cell cycle, division, growth and fate | B( .027) | 1.08 | 0.92 | BQ-downreg |
| 12 | 117768545 | XP_034452902.1 | OSTF1 | Osteoclast stimulating factor 1 | Cell cycle, division, growth and fate | B( .014) | 1.09 | 0.92 | BQ-downreg |
| 13 | 117765831 | XP_034448251.1 | MTAP | S-methyl-5'-thioadenosine phosphorylase | Translation | B( .004) | 1.09 | 0.92 | BQ-downreg |
| 14 | 117766769 | XP_034450010.1 | JPT2 | Jupiter microtubule associated homolog 2 | Cell cycle, division, growth and fate | B( .013) | 1.09 | 0.92 | BQ-downreg |
| 15 | 117771385 | XP_034457559.1 | RPL17 | 60S ribosomal protein L17 | Translation | B( .023) | 1.09 | 0.92 | BQ-downreg |
| 16 | 117767555 | XP_034451214.1 | LSM6 | LSM6 homolog, U6 small nuclear RNA and mRNA degradation associated | Transcription | B( .009) | 1.10 | 0.91 | BQ-downreg |
| 17 | 117769845 | XP_034454897.1 | PPID | Peptidylprolyl isomerase D | Protein folding | B( .006) | 1.10 | 0.91 | BQ-downreg |
| 18 | 117761009 | XP_034440436.1 | IPO9 | Importin 9 | Protein transport | B( .016) | 1.10 | 0.91 | BQ-downreg |
| 19 | 117767049 | XP_034450480.1 | PHB | Prohibitin | Mitochondrial biogenesis | B( .036) | 1.11 | 0.90 | BQ-downreg |
| 20 | 117753795 | XP_034428030.1 | PDIA4 | Protein disulfide isomerase family A member 4 | Protein folding | B( .041) | 1.11 | 0.90 | BQ-downreg |
| 21 | 117759425 | XP_034437442.1 | RBBP7 | Histone-binding protein RBBP7 | Cell cycle, division, growth and fate | B( .042) | 1.11 | 0.90 | BQ-downreg |
| 22 | 117764281 | XP_034445816.1 | NUDT16 | U8 snoRNA-decapping enzyme | Cell cycle, division, growth and fate | B( .023) | 1.11 | 0.90 | BQ-downreg |
| 23 | 117774105 | XP_034462107.1 | C1QBP | Complement component 1 Q subcomponent-binding protein, mitochondrial | Translation | B( .017) | 1.11 | 0.90 | BQ-downreg |
| 24 | 117761960 | XP_034442253.1 | ENO1 | Enolase 1 | Energy metabolism | B( .018) | 1.11 | 0.90 | BQ-downreg |
| 25 | 117760081 | XP_034438718.1 | DNAJC19 | Mitochondrial import inner membrane translocase subunit TIM14 | Protein folding | B( .010) | 1.12 | 0.89 | BQ-downreg |
| 26 | 117774668 | XP_034463139.1 | TUBA | Tubulin alpha chain | Cell cycle, division, growth and fate | B( .037) | 1.12 | 0.89 | BQ-downreg |
| 27 | 117764430 | XP_034446129.1 | B3GALT2 | Beta-1,3-galactosyltransferase 5-like | Protein folding | B( .017) | 1.13 | 0.88 | BQ-downreg |
| 28 | 117755614 | XP_034431437.1 | PCCB | Propionyl-CoA carboxylase beta chain, mitochondrial | Energy metabolism | B( .012) | 1.13 | 0.88 | BQ-downreg |
| 29 | 117760044 | XP_034438635.1 | HMCN1 | Hemicentin 1 | Cell cycle, division, growth and fate | B( .021) | 1.13 | 0.88 | BQ-downreg |
| 30 | 117767010 | XP_034450409.1 | SNRNP70 | Small nuclear ribonucleoprotein U1 subunit 70 kDa | Transcription | B( .027) | 1.13 | 0.88 | BQ-downreg |
| 31 | 117769084 | XP_034453583.1 | ACSL4 | Acyl-CoA synthetase long chain family member 4 | Lipid metabolism | B( .035) | 1.14 | 0.88 | BQ-downreg |
| 32 | 117762830 | XP_034443368.1 | DNAJA4 | DnaJ heat shock protein family (Hsp40) member A4 | Protein folding | B( .012) | 1.14 | 0.87 | BQ-downreg |
| 33 | 117776536 | XP_034466413.1 | MECR | Enoyl-[acyl-carrier-protein] reductase, mitochondrial | Lipid metabolism | B( .041) | 1.15 | 0.87 | BQ-downreg |
| 34 | 117775309 | XP_034464242.1 | PGAM1 | Phosphoglycerate mutase 1 | Energy metabolism | B( .021) | 1.15 | 0.87 | BQ-downreg |
| 35 | 117770240 | XP_034455309.1 | DDX6 | ATP-dependent RNA helicase DDX6 | Cell cycle, division, growth and fate | B( .028) | 1.15 | 0.87 | BQ-downreg |
| 36 | 117778281 | XP_034469638.1 | RANBP3 | RAN binding protein 3 | Protein transport | B( .022) | 1.15 | 0.87 | BQ-downreg |
| 37 | 117752505 | XP_034425746.1 | DCXR | Dicarbonyl and L-xylulose reductase | Energy metabolism | B( .037) | 1.16 | 0.86 | BQ-downreg |
| 38 | 117776903 | XP_034467184.1 | BCKDHB | 2-oxoisovalerate dehydrogenase subunit beta, mitochondrial | Energy metabolism | B( .018) | 1.17 | 0.86 | BQ-downreg |
| 39 | 117766179 | XP_034448855.1 | FTH1 | Ferritin, middle subunit isoform X1 | Others | B( .001) | 1.17 | 0.86 | BQ-downreg |
| 40 | 117760188 | XP_034438911.1 | QSOX1 | Quiescin sulfhydryl oxidase 1 | Protein folding | B( .005) | 1.17 | 0.85 | BQ-downreg |
| 41 | 117768165 | XP_034452215.1 | SMARCAD1 | SWI/SNF-related, matrix-associated actin-dependent regulator of chromatin | Cell cycle, division, growth and fate | B( .015) | 1.18 | 0.85 | BQ-downreg |
| 42 | 117774762 | XP_034463285.1 | CKM | Creatine kinase, M-type | Energy metabolism | B( .035) | 1.19 | 0.84 | BQ-downreg |
| 43 | 117758658 | XP_034436395.1 | MTIF3 | Translation initiation factor IF-3, mitochondrial | Translation | B( .017) | 1.19 | 0.84 | BQ-downreg |
| 44 | 117774907 | XP_034463503.1 | DHRS9 | Dehydrogenase/reductase SDR family member 9 | Metabolism of cofactors and vitamins | B( .006) | 1.20 | 0.83 | BQ-downreg |
| 45 | 117755535 | XP_034431292.1 | PTTG1IP | Pituitary tumor-transforming gene 1 protein-interacting protein | Protein transport | B( .035) | 1.21 | 0.83 | BQ-downreg |
| 46 | 117759696 | XP_034437849.1 | MAGOH | Mago homolog, exon junction complex subunit | Transcription | B( .035) | 1.21 | 0.83 | BQ-downreg |
| 47 | 117767482 | XP_034451051.1 | HIP1R | Huntingtin-interacting protein 1-related protein | Protein transport | B( .015) | 1.22 | 0.82 | BQ-downreg |
| 48 | 117765842 | XP_034448270.1 | GCDH | Glutaryl-CoA dehydrogenase, mitochondrial | Lipid metabolism | B( .014) | 1.23 | 0.82 | BQ-downreg |
| 49 | 117770565 | XP_034456028.1 | CAP1 | Adenylyl cyclase-associated protein 1 | Cell cycle, division, growth and fate | B( .027) | 1.23 | 0.81 | BQ-downreg |
| 50 | 117764881 | XP_034446894.1 | KIF1B | Kinesin family member 1B | Cell cycle, division, growth and fate | B( .047) | 1.24 | 0.81 | BQ-downreg |
| 51 | 117762890 | XP_034443496.1 | CNOT1 | CCR4-NOT transcription complex subunit 1 | Protein degradation and synthesis inhibition | B( .044) | 1.25 | 0.80 | BQ-downreg |
| 52 | 117764636 | XP_034446468.1 | LIMA1 | LIM domain and actin binding 1 | Cell cycle, division, growth and fate | B( .008) | 1.25 | 0.80 | BQ-downreg |
| 53 | 117758633 | XP_034436360.1 | PPIL4 | Peptidylprolyl isomerase like 4 | Protein folding | B( .013) | 1.26 | 0.79 | BQ-downreg |
| 54 | 117770713 | XP_034456270.1 | PBXIP1 | PBX homeobox interacting protein 1 | Cell cycle, division, growth and fate | B( .028) | 1.27 | 0.79 | BQ-downreg |
| 55 | 117755249 | XP_034430776.1 | GATD3A | Glutamine amidotransferase like class 1 domain containing 3A, mitochondrial | Mitochondrial biogenesis | B( .000) | 1.27 | 0.79 | BQ-downreg |
| 56 | 117775505 | XP_034464599.1 | MCFD2 | Multiple coagulation factor deficiency 2, ER cargo receptor complex subunit | Protein transport | B( .018) | 1.28 | 0.78 | BQ-downreg |
| 57 | 117772522 | XP_034459609.1 | GFM1 | G elongation factor mitochondrial 1 | Translation | B( .029) | 1.29 | 0.78 | BQ-downreg |
| 58 | 117755704 | XP_034431608.1 | RBBP4 | Histone-binding protein RBBP4 | Cell cycle, division, growth and fate | B( .042) | 1.31 | 0.76 | BQ-downreg |
| 59 | 5405699 | YP_001403139.1 | MT-ND5 | NADH-ubiquinone oxidoreductase chain 5 | Mitochondrial biogenesis | B( .005) | 1.32 | 0.76 | BQ-downreg |
| 60 | 117775592 | XP_034464795.1 | EIF3L | Eukaryotic translation initiation factor 3 subunit L | Translation | B( .017) | 1.33 | 0.75 | BQ-downreg |
| 61 | 117766124 | XP_034448738.1 | LITAF | Lipopolysaccharide-induced tumor necrosis factor-alpha factor homolog | Protein degradation and synthesis inhibition | B( .023) | 1.34 | 0.75 | BQ-downreg |
| 62 | 117768498 | XP_034452796.1 | TCN2 | Transcobalamin 2 | Metabolism of cofactors and vitamins | B( .013) | 1.54 | 0.65 | BQ-downreg |
| 63 | 117767343 | XP_034450787.1 | COQ7 | 5-demethoxyubiquinone hydroxylase, mitochondrial | Lipid metabolism | B( .033) | 1.67 | 0.60 | BQ-downreg |
| 64 | 117774510 | XP_034462875.1 | SOD1 | Superoxide dismutase 1 | Redox/Detox related | B( .012) | 1.85 | 0.54 | BQ-downreg |

**Table S3. STRING protein-protein interaction networks enrichment analysis.** Network enrichment analyses for differentially regulated proteins (down-regulated in poor quality eggs, N = 64 and up-regulated in poor quality eggs) which were resolved in a STRING subnetworks. Gene Ontology (GO) Biological processes, molecular functions and cellular components, along with KEGG and Reactome pathways are given. Proteins were mapped against human database. GO-term IDs, description of processes and pathways, number of query proteins mapping to the total number of representative proteins present in the database and the false discovery rates (FDR) are given in each column. PPI enrichment $p$ values indicating the existence of non-random nodes and therefore the significance of the observed edge numbers are given in the header panels.

**Network Statistics for proteins down-regulated in poor quality eggs N = 64**

| | |
|---|---|
| Reference database: | Human |
| number of nodes: | 62 |
| number of edges: | 65 |
| expected number of edges: | 28 |
| average node degree: | 2.1 |
| avg. local clustering coefficient: | 0.316 |
| PPI enrichment p-value: | 1.70E-09 |

**Biological Process (GO)**

| *GO-term* | *Description* | *#Protein* | *FDR* |
|---|---|---|---|
| GO:1901575 | organic substance catabolic process | 17 of 1609 | 0.0019 |
| GO:1901566 | organonitrogen compound biosynthetic process | 16 of 1370 | 0.0019 |
| GO:1901564 | organonitrogen compound metabolic process | 34 of 5281 | 0.0019 |
| GO:0055114 | oxidation-reduction process | 14 of 932 | 0.0019 |
| GO:0055086 | nucleobase-containing small molecule metabolic process | 11 of 662 | 0.0019 |
| GO:0051186 | cofactor metabolic process | 10 of 467 | 0.0019 |
| GO:0044282 | small molecule catabolic process | 9 of 388 | 0.0019 |
| GO:0044281 | small molecule metabolic process | 19 of 1779 | 0.0019 |
| GO:0044248 | cellular catabolic process | 18 of 1646 | 0.0019 |
| GO:0043603 | cellular amide metabolic process | 12 of 732 | 0.0019 |
| GO:0034655 | nucleobase-containing compound catabolic process | 9 of 394 | 0.0019 |
| GO:0019752 | carboxylic acid metabolic process | 13 of 854 | 0.0019 |
| GO:0008152 | metabolic process | 48 of 9569 | 0.0019 |
| GO:0006732 | coenzyme metabolic process | 8 of 297 | 0.0019 |
| GO:0043604 | amide biosynthetic process | 9 of 495 | 0.0021 |
| GO:0017144 | drug metabolic process | 10 of 622 | 0.0021 |
| GO:1901135 | carbohydrate derivative metabolic process | 13 of 1083 | 0.0023 |
| GO:0044237 | cellular metabolic process | 44 of 8797 | 0.0023 |
| GO:0009066 | aspartate family amino acid metabolic process | 4 of 55 | 0.0024 |
| GO:0016071 | mRNA metabolic process | 10 of 667 | 0.0031 |
| GO:0034641 | cellular nitrogen compound metabolic process | 31 of 5126 | 0.0034 |
| GO:0006402 | mRNA catabolic process | 6 of 207 | 0.0034 |
| GO:0051188 | cofactor biosynthetic process | 6 of 218 | 0.0044 |
| GO:0009117 | nucleotide metabolic process | 9 of 576 | 0.0049 |
| GO:0046395 | carboxylic acid catabolic process | 6 of 237 | 0.0062 |
| GO:0034404 | nucleobase-containing small molecule biosynthetic process | 5 of 149 | 0.0062 |
| GO:0072521 | purine-containing compound metabolic process | 8 of 478 | 0.0065 |
| GO:0046434 | organophosphate catabolic process | 5 of 152 | 0.0065 |
| GO:0032787 | monocarboxylic acid metabolic process | 8 of 477 | 0.0065 |
| GO:0009987 | cellular process | 58 of 14652 | 0.0065 |
| GO:1901137 | carbohydrate derivative biosynthetic process | 9 of 625 | 0.0068 |
| GO:0006810 | transport | 26 of 4130 | 0.0074 |
| GO:0006913 | nucleocytoplasmic transport | 6 of 263 | 0.0083 |
| GO:0034248 | regulation of cellular amide metabolic process | 7 of 379 | 0.0084 |
| GO:0009108 | coenzyme biosynthetic process | 5 of 167 | 0.0084 |
| GO:0071704 | organic substance metabolic process | 43 of 9135 | 0.0099 |
| GO:0019637 | organophosphate metabolic process | 11 of 1011 | 0.0115 |
| GO:0009204 | deoxyribonucleoside triphosphate catabolic process | 2 of 7 | 0.012 |
| GO:0009166 | nucleotide catabolic process | 4 of 101 | 0.012 |
| GO:0000956 | nuclear-transcribed mRNA catabolic process | 5 of 191 | 0.013 |
| GO:0072329 | monocarboxylic acid catabolic process | 4 of 106 | 0.0135 |
| GO:0044283 | small molecule biosynthetic process | 8 of 569 | 0.0141 |
| GO:0006807 | nitrogen compound metabolic process | 40 of 8349 | 0.0141 |
| GO:0006520 | cellular amino acid metabolic process | 6 of 308 | 0.0143 |
| GO:0072522 | purine-containing compound biosynthetic process | 5 of 206 | 0.0161 |
| GO:0050684 | regulation of mRNA processing | 4 of 114 | 0.0161 |
| GO:1901605 | alpha-amino acid metabolic process | 5 of 209 | 0.0168 |
| GO:0009058 | biosynthetic process | 27 of 4714 | 0.0177 |

| | | | |
|---|---|---|---|
| GO:0006554 | lysine catabolic process | 2 of 10 | 0.018 |
| GO:0051170 | import into nucleus | 4 of 122 | 0.0189 |
| GO:0006733 | oxidoreduction coenzyme metabolic process | 4 of 124 | 0.0198 |
| GO:0006790 | sulfur compound metabolic process | 6 of 343 | 0.0215 |
| GO:0016575 | histone deacetylation | 3 of 57 | 0.0234 |
| GO:0009179 | purine ribonucleoside diphosphate metabolic process | 3 of 58 | 0.0242 |
| GO:1903311 | regulation of mRNA metabolic process | 5 of 238 | 0.0246 |
| GO:0046365 | monosaccharide catabolic process | 3 of 59 | 0.0246 |
| GO:0006518 | peptide metabolic process | 7 of 497 | 0.0246 |
| GO:0000288 | nuclear-transcribed mRNA catabolic process, deadenylation-dependent decay | 3 of 59 | 0.0246 |
| GO:0009141 | nucleoside triphosphate metabolic process | 5 of 246 | 0.0268 |
| GO:1901576 | organic substance biosynthetic process | 26 of 4656 | 0.028 |
| GO:0008090 | retrograde axonal transport | 2 of 16 | 0.0324 |
| GO:0045055 | regulated exocytosis | 8 of 691 | 0.0333 |
| GO:0044238 | primary metabolic process | 40 of 8808 | 0.0335 |
| GO:0010629 | negative regulation of gene expression | 13 of 1670 | 0.0411 |
| GO:0009264 | deoxyribonucleotide catabolic process | 2 of 19 | 0.0411 |
| GO:0048024 | regulation of mRNA splicing, via spliceosome | 3 of 77 | 0.042 |
| GO:0044242 | cellular lipid catabolic process | 4 of 167 | 0.042 |
| GO:0046386 | deoxyribose phosphate catabolic process | 2 of 20 | 0.0431 |
| GO:0044249 | cellular biosynthetic process | 25 of 4567 | 0.0431 |
| GO:0006555 | methionine metabolic process | 2 of 20 | 0.0431 |
| GO:0090407 | organophosphate biosynthetic process | 7 of 577 | 0.0446 |
| GO:0009165 | nucleotide biosynthetic process | 5 of 291 | 0.0446 |
| GO:0009150 | purine ribonucleotide metabolic process | 6 of 425 | 0.0446 |
| GO:0009062 | fatty acid catabolic process | 3 of 84 | 0.0473 |

**Molecular Function (GO)**

| ***GO-term*** | ***Description*** | ***#Protein*** | ***FDR*** |
|---|---|---|---|
| GO:0016491 | oxidoreductase activity | 12 of 716 | 0.00077 |
| GO:0016853 | isomerase activity | 6 of 147 | 0.0014 |
| GO:0043022 | ribosome binding | 4 of 50 | 0.0027 |
| GO:0043021 | ribonucleoprotein complex binding | 5 of 116 | 0.0032 |
| GO:0003824 | catalytic activity | 33 of 5592 | 0.0032 |
| GO:0030619 | U1 snRNA binding | 2 of 7 | 0.0183 |
| GO:0047429 | nucleoside-triphosphate diphosphatase activity | 2 of 10 | 0.0287 |
| GO:0001846 | opsonin binding | 2 of 14 | 0.0452 |

**Cellular Component (GO)**

| ***GO-term*** | ***Description*** | ***#Protein*** | ***FDR*** |
|---|---|---|---|
| GO:0005737 | cytoplasm | 60 of 11238 | 4.26E-10 |
| GO:0005739 | mitochondrion | 20 of 1531 | 4.58E-06 |
| GO:0043231 | intracellular membrane-bounded organelle | 53 of 10365 | 7.24E-06 |
| GO:0005622 | intracellular | 61 of 14286 | 7.24E-06 |
| GO:0043229 | intracellular organelle | 57 of 12193 | 7.30E-06 |
| GO:0070013 | intracellular organelle lumen | 36 of 5162 | 7.86E-06 |
| GO:0043227 | membrane-bounded organelle | 54 of 11244 | 1.49E-05 |
| GO:0005759 | mitochondrial matrix | 9 of 463 | 0.00047 |
| GO:0005623 | cell | 61 of 16271 | 0.0039 |
| GO:0005829 | cytosol | 29 of 4958 | 0.0053 |
| GO:1904724 | tertiary granule lumen | 3 of 55 | 0.0184 |
| GO:0120111 | neuron projection cytoplasm | 3 of 71 | 0.0347 |
| GO:0099568 | cytoplasmic region | 6 of 402 | 0.0351 |
| GO:0005740 | mitochondrial envelope | 8 of 722 | 0.0373 |
| GO:0120114 | Sm-like protein family complex | 3 of 78 | 0.0375 |
| GO:0000932 | P-body | 3 of 81 | 0.0394 |

**KEGG Pathways**

| ***Pathway*** | ***Description*** | ***#Protein*** | ***FDR*** |
|---|---|---|---|
| hsa01100 | Metabolic pathways | 19 of 1250 | 6.13E-07 |
| hsa03018 | RNA degradation | 5 of 77 | 0.00028 |
| hsa00280 | Valine, leucine and isoleucine degradation | 3 of 48 | 0.0138 |
| hsa00071 | Fatty acid degradation | 3 of 44 | 0.0138 |
| hsa04217 | Necroptosis | 4 of 155 | 0.0277 |
| hsa00010 | Glycolysis / Gluconeogenesis | 3 of 68 | 0.0277 |

**Reactome Pathways**

| Pathway | Description | #Protein | FDR |
|---|---|---|---|
| HSA-1430728 | Metabolism | 19 of 2032 | 0.0033 |

1298

**Network Statistics for proteins up-regulated in poor quality eggs N = 51**

| | |
|---|---|
| Reference database: | Human |
| Number of nodes: | 50 |
| Number of edges: | 36 |
| Expected number of edges: | 20 |
| Average node degree: | 1.44 |
| Avg. local clustering coefficient: | 0.405 |
| PPI enrichment p-value: | 0.000574 |

**Biological Process (GO)**

| GO-term | Description | #Protein | FDR |
|---|---|---|---|
| GO:0071704 | organic substance metabolic process | 39 of 9135 | 0.009 |
| GO:0044238 | primary metabolic process | 38 of 8808 | 0.009 |
| GO:0044237 | cellular metabolic process | 38 of 8797 | 0.009 |
| GO:0009987 | cellular process | 49 of 14652 | 0.009 |
| GO:0008152 | metabolic process | 40 of 9569 | 0.009 |
| GO:0044403 | symbiont process | 9 of 650 | 0.0091 |
| GO:0008380 | RNA splicing | 7 of 391 | 0.0131 |
| GO:0006091 | generation of precursor metabolites and energy | 7 of 400 | 0.0132 |
| GO:0016032 | viral process | 8 of 571 | 0.0144 |
| GO:0007005 | mitochondrion organization | 7 of 424 | 0.0144 |
| GO:0006839 | mitochondrial transport | 5 of 223 | 0.034 |

**Cellular Component (GO)**

| GO-term | Description | #Protein | FDR |
|---|---|---|---|
| GO:0005750 | mitochondrial respiratory chain complex III | 3 of 13 | 0.002 |
| GO:0005743 | mitochondrial inner membrane | 8 of 456 | 0.002 |
| GO:0043231 | intracellular membrane-bounded organelle | 40 of 10365 | 0.0027 |
| GO:0031966 | mitochondrial membrane | 9 of 679 | 0.0027 |
| GO:0005737 | cytoplasm | 42 of 11238 | 0.0027 |
| GO:0031967 | organelle envelope | 11 of 1146 | 0.0029 |
| GO:0005829 | cytosol | 25 of 4958 | 0.003 |
| GO:1904813 | ficolin-1-rich granule lumen | 4 of 125 | 0.0057 |
| GO:0098800 | inner mitochondrial membrane protein complex | 4 of 128 | 0.0058 |
| GO:0005739 | mitochondrion | 12 of 1531 | 0.006 |
| GO:0070013 | intracellular organelle lumen | 24 of 5162 | 0.01 |
| GO:0043229 | intracellular organelle | 42 of 12193 | 0.01 |
| GO:0032991 | protein-containing complex | 23 of 4792 | 0.01 |
| GO:0005622 | intracellular | 46 of 14286 | 0.01 |
| GO:0071013 | catalytic step 2 spliceosome | 3 of 99 | 0.0183 |
| GO:1902494 | catalytic complex | 9 of 1295 | 0.0375 |
| GO:0071011 | precatalytic spliceosome | 2 of 41 | 0.0375 |
| GO:0005634 | nucleus | 27 of 6892 | 0.0375 |

**Local STRING network cluster**

| Cluster | Description | #Protein | FDR |
|---|---|---|---|
| CL:15986 | mRNA Splicing | 7 of 185 | 0.00013 |
| CL:22476 | mitochondrial respiratory chain complex III | 3 of 6 | 0.00016 |
| CL:22328 | respirasome | 5 of 94 | 0.00025 |
| CL:22327 | Oxidative phosphorylation | 6 of 153 | 0.00025 |
| CL:15989 | mRNA Splicing - Major Pathway | 6 of 157 | 0.00025 |
| CL:15992 | mRNA Splicing - Major Pathway | 5 of 146 | 0.0012 |
| CL:22417 | Complex I biogenesis | 2 of 11 | 0.009 |
| CL:15994 | mRNA Splicing - Major Pathway | 4 of 140 | 0.009 |
| CL:13515 | protein neddylation | 2 of 11 | 0.009 |

**KEGG Pathways**

| Pathway | Description | #Protein | FDR |
|---|---|---|---|
| hsa05010 | Alzheimer's disease | 5 of 168 | 0.0041 |
| hsa05012 | Parkinson's disease | 4 of 142 | 0.0072 |
| hsa04714 | Thermogenesis | 5 of 228 | 0.0072 |
| hsa01100 | Metabolic pathways | 11 of 1250 | 0.0072 |
| hsa00190 | Oxidative phosphorylation | 4 of 131 | 0.0072 |
| hsa04260 | Cardiac muscle contraction | 3 of 76 | 0.0095 |
| hsa05016 | Huntington's disease | 4 of 193 | 0.012 |
| hsa00020 | Citrate cycle (TCA cycle) | 2 of 30 | 0.0199 |

| | | | |
|---|---|---|---|
| hsa03040 | Spliceosome | 3 of 130 | 0.0279 |
| hsa04932 | Non-alcoholic fatty liver disease (NAFLD) | 3 of 149 | 0.0363 |
| **Reactome Pathways** | | | |
| ***Pathway*** | ***Description*** | ***#Protein*** | ***FDR*** |
| HSA-1428517 | The citric acid (TCA) cycle and respiratory electron transport | 7 of 173 | 0.00011 |
| HSA-163200 | Respiratory electron transport, ATP synthesis by chemiosmotic coupling, and heat production by uncoupling proteins. | 6 of 123 | 0.00014 |
| HSA-611105 | Respiratory electron transport | 5 of 100 | 0.00071 |
| HSA-72203 | Processing of Capped Intron-Containing Pre-mRNA | 6 of 234 | 0.0024 |
| HSA-72163 | mRNA Splicing - Major Pathway | 5 of 178 | 0.0061 |
| HSA-1430728 | Metabolism | 14 of 2032 | 0.0187 |
| HSA-1169408 | ISG15 antiviral mechanism | 3 of 69 | 0.0311 |
| HSA-8953854 | Metabolism of RNA | 7 of 652 | 0.0403 |
| **INTERPRO Protein Domains and Features** | | | |
| ***Domain*** | ***Description*** | ***#Protein*** | ***FDR*** |
| IPR016024 | Armadillo-type fold | 7 of 333 | 0.0045 |
| IPR011989 | Armadillo-like helical | 5 of 183 | 0.011 |

**Table S4. List of primers and probes used for relative quantification of gene expression and mtDNA quantification using TaqMan based qPCR.** Names, sequences, length, GC content and annealing temperatures of primers and probes along with expected amplicon size in base pairs (bp) are given in each column.

| Name | Sequence | Length | GC | Tm | Amplicon (bp) |
|---|---|---|---|---|---|
| *18S*_Forward | TTGGCGAGGACCTGACTGTATTTCCCG | 27 | 55.6 | 62.1 | 146 |
| *18S*_Reverse | ACCAAGAGGGCAGGAGAACTCACTGAGG | 28 | 57.1 | 62.8 | |
| *18S*_Probe | TGCATGATGGTCACCACCCGCTCCACCT | 28 | 60.7 | 66.8 | |
| *mt-nd5*_Forward | GGCTTGAATAGCAGCAAACCTAAACTCCTGG | 31 | 48.4 | 61.4 | 154 |
| *mt-nd5*_Reverse | AGCGGAGGGTAGTCAAGGGTGAAGTCC | 27 | 59.3 | 62.9 | |
| *mt-nd5*_Probe | ACTCATCATTGCAGCAACTGGCAAATCGGCTC | 32 | 50 | 64.8 | |
| *cap1*_Forward | TTGTAAGAAGTTGGGGCTGGTGTTTGACC | 29 | 48.3 | 61.1 | 144 |
| *cap1*_Reverse | TGGCTCAGGTAAATGTGGCAACCGTC | 26 | 53.8 | 61.4 | |
| *cap1*_Probe | CGTCGTCGGCATCGTGGAGATCATCAACTGC | 31 | 58.1 | 65.9 | |
| *dhrs9*_Forward | AGGGAGAAGACGATCTGAAGAAGTCCTGC | 29 | 51.7 | 61.1 | 139 |
| *dhrs9*_Reverse | ACTACAGCCCATAAACCTCGCTCGCC | 26 | 57.7 | 62.4 | |
| *dhrs9*_Probe | CCAGCAGCCTGACCACCATGCACCTCGAT | 29 | 62.1 | 67.2 | |
| *gcn1*_Forward | TGTGCTGAAGTAACACACTCCGCAGCTC | 28 | 53.6 | 62.8 | 150 |
| *gcn1*_Reverse | ACCCAACCATCCGCAAGAACATTACCACC | 29 | 51.7 | 62.7 | |
| *gcn1*_Probe | AGTTCACCAACACAGCCCGCTGACGCC | 27 | 63 | 66.8 | |
| *ghitm*_Forward | ATGTGTGCCCCAAGCGAGAAGTTCC | 25 | 56 | 61.4 | 137 |
| *ghitm*_Reverse | GCCACTGAGTAAAGACCTGCTCCAAATGC | 29 | 51.7 | 61.7 | |
| *ghitm*_Probe | TCGGATCAATGTTCCTGCCGCCCAGCTC | 28 | 60.7 | 65.9 | |
| *uqcrfs1*_Forward | ACATCAGAGCGACGATAGTCTCCAAAGTCAG | 31 | 48.4 | 61.3 | 142 |
| *uqcrfs1*_Reverse | TGCAGGCAGCGAAACACTCCGTTAAG | 26 | 53.8 | 61.8 | |
| *uqcrfs1*_Probe | AGCAGCAGCGGGCACCAGCTCCTT | 24 | 66.7 | 66.8 | |
| *gatd3a*_Forward | GCTCAGCAGGCAACCACCTCTTCTTC | 26 | 57.7 | 61.6 | 138 |
| *gatd3a*_Reverse | CGCATTCATGTGCAACGCTCCCATC | 25 | 56 | 61.4 | |
| *gatd3a*_Probe | TCAAGCCAGCTTCAGCACGTCCCGCA | 26 | 61.5 | 66.4 | |
| *fbp1a*_Forward | TGCTTCGATCCTTTGGACGGCTCCTC | 26 | 57.7 | 62.4 | 101 |
| *fbp1a*_Reverse | TCACACGGCTCATCGTCTGTGGTCTTTC | 28 | 53.6 | 62.4 | |
| *fbp1a*_Probe | TCGACTGTCTCGTCTCCATCGGCACCA | 27 | 59.3 | 64.5 | |
| *atp5f1b*_Forward | GACCCACCATGTAGAACGCTTGCTCC | 26 | 57.7 | 61.5 | 148 |
| *atp5f1b*_Reverse | TTCTTGTCGCAGCCTTTCCAGGTCG | 25 | 56 | 61.4 | |
| *atp5f1b*_Probe | TGTTTCCTTGAGAGGCACCAGCTTGCCCAT | 30 | 53.3 | 65.3 | |
| *acylb*_Forward | TGTCGTCCACTGCAACCATCAGCCTC | 26 | 57.7 | 62.8 | 153 |
| *acylb*_Reverse | AGCCGTTTGTTCCACACACACAGGAAG | 27 | 51.9 | 61.6 | |
| *acylb*_Probe | CGTCCACGTCGCCCACCTCCACTCCT | 26 | 69.2 | 67.2 | |
| *gcdh*_Forward | GAGCTGATCTGGCTTCACCACGAGAC | 26 | 57.7 | 61 | 117 |
| *gcdh*_Reverse | GCCGTGCAGAATCGCTTTTCCTATGC | 26 | 53.8 | 61.1 | |
| *gcdh*_Probe | TGCTCCTGCGTGAGGCTGTCCCAGTCA | 27 | 63 | 66.7 | |
| *mt-atp6*_Forward | GCCCTCATCCCTGTACTTATT | 21 | 47.6 | 57.9 | 94 |
| *mt-atp6*_Reverse | CTGCTGTGAGATTTGCTGTTAG | 22 | 45.5 | 58.4 | |
| *mt-atp6*_Probe | TCATTCGACCCTTAGCTCTCGGTGT | 25 | 64.6 | 52 | |
| *uqcrb*_Forward | TTAACATGAAGCAGCAGATCC | 21 | 42.9 | 50.5 | 112 |
| *uqcrb*_Reverse | TCTATTTCTTTACGCTCACGG | 21 | 42.9 | 50.1 | |
| *uqcrb*_Probe | AGGATATCAGCTACCTGTCACCATATCTGAA | 31 | 41.9 | 58.3 | |
| *cyc1*_Forward | AACATCTCAGTAAGTCGTCTGAAGACCC | 28 | 46.4 | 58 | 133 |
| *cyc1*_Reverse | CAACCTTGTGCTTTCCACCATTCTCC | 26 | 50 | 58.5 | |
| *cyc1*_Probe | CATTTGTCCAGAAGTGTGCCCAGTGCC | 27 | 55.6 | 62.2 | |
| *fh*_Forward | AGCATTTCACTTGGCTTCACCCAC | 24 | 50 | 58.1 | 105 |
| *fh*_Reverse | CAAGACGGCTCACAAGAAAGGCAC | 24 | 54.2 | 58.7 | |
| *fh*_Probe | TGCTCCTCGGTCACGTATCCCAGCTCAA | 28 | 57.1 | 64.4 | |
| *pdia4*_Forward | CGTCCAAACCCCAACACTTTTCCTC | 25 | 52 | 58.4 | 100 |
| *pdia4*_Reverse | TCCATCCCTCAGTACCCTCTGTACTC | 26 | 53.8 | 58.1 | |
| *pdia4*_Probe | TCCAGGTAGTAATTGATCTGCACGGCAGCCA | 31 | 51.6 | 64.4 | |
| *ppid*_Forward | CGCAGTTCTTCATCACCACTGTCC | 24 | 54.2 | 58 | 101 |
| *ppid*_Reverse | CTCCAGCATTTTTACGACACCCATTCC | 27 | 48.1 | 58.5 | |
| *ppid*_Probe | CCCTCACTTAGATGATAAGCACGTTGTCTTCGG | 33 | 48.5 | 62 | |
| *phb*_Forward | GACGGGAACGGCAATCAAAGATGATAG | 27 | 48.1 | 58.2 | 102 |
| *phb*_Reverse | TCGACAGGTTCAGAGGAGTGCAAG | 24 | 54.2 | 58.4 | |
| *phb*_Probe | TGATGAAGTGAGTCCCTTCACCAGCAACAGT | 31 | 48.4 | 62.9 | |
| *vdac*_Forward | CTGTACCTCCCACCTACATTGACCTTG | 27 | 51.9 | 58.4 | 116 |
| *vdac*_Reverse | CTCCAGTCCATTGTCCGACTTTGTTTTC | 28 | 46.4 | 58.5 | |
| *vdac*_Probe | TGGCTTCGGGCTCATCAAGCTGGACC | 26 | 61.5 | 64.5 | |
| *mecr*_Forward | CAGGATTTTAACCAGGACATCCTTTGCAC | 29 | 44.8 | 58.6 | 107 |
| *mecr*_Reverse | GCTCAGCCCTTCAGTACAGACAACAC | 26 | 53.8 | 59.5 | |
| *mecr*_Probe | TGGGTGGCAAATCTATATCTTCCAACTGGACGAC | 34 | 47.1 | 62.8 | |
| *gfm1*_Forward | TCTCACCTGTCCAACAATCTCTGTTTCC | 28 | 46.4 | 58.6 | 106 |
| *gfm1*_Reverse | TAATGTGACGATCAGACCTGAAGCTGCC | 28 | 50 | 60.5 | |
| *gfm1*_Probe | TGAGCCAACACGTCGTCCGCTGAG | 24 | 62.5 | 63.2 | |
| *sod1*_Forward | ACACAAATGGGTGCATGAGTGCAG | 24 | 50 | 58.4 | 107 |
| *sod1*_Reverse | AGTCACATTCCCCAGGTCTCCAAC | 24 | 54.2 | 58.1 | |
| *sod1*_Probe | TGCTGGTCCTACTGATGCAGACAGGCAC | 28 | 57.1 | 63.7 | |

**Table S5. List of reference peptides used for PRM based LC-MS/MS.** Names, amino acid sequences, length in amino acids (aa), charge, peptide spectral matches (PSMs), miscleavage potentials, significant differences ($p < 0.05$) in peptide abundance between good and poor quality eggs (based on TMT labeling based LC-MS/MS), suitability for PRM and hydrophobicity rates (%) (based on Peptide Synthesis and Proteotypic Peptide Analysis, ThermoFisher Scientific) are given in each column. See the **Material and Methods** section for details on selection of reference peptides used in this study.

| Protein | Peptide | Length (aa) | Charge | PSMs | Miscleavage | $p < 0.05$ | Suitablity for PRM | Hydrophobicty (%) |
|---|---|---|---|---|---|---|---|---|
| DHRS9_5 | NVETLWA(K) | 8 | 2 | 2 | 0 | 0.043 | YES | 25.49 |
| DHRS9_6 | VVNVASVFG( R) | 10 | 2 | 2 | 0 | 0.027 | YES | 25.11 |
| DHRS9_7 | YSPGWDA(K) | 8 | 2 | 2 | 0 | 0.002 | YES | 18.96 |
| GATD3A_1 | DDKYPDTTGTAEAINQLGC(K) | 20 | 3 | 2 | 0 | 0.005 | YES | 25.18 |
| GATD3A_2 | GDIQDLS(K) | 8 | 2 | 2 | 0 | 0.007 | YES | 14.9 |
| GATD3A_3 | NVLVESA(R) | 8 | 2 | 2 | 0 | - | YES | 16.73 |
| MT-ND5_1 | AIEAANRPLVSSTSNIQ( R) | 18 | 3 | 2 | 0 | 0.103 | YES | 27.25 |
| MT-ND5_2 | FSPLSPINENNPAVINPLK( R) | 20 | 3 or 4 | 4 | 1 | - | YES | 36.65 |
| MT-ND5_3 | FSPLSPINENNPAVINPL(K) | 19 | 3 or 4 | 4 | 0 | - | YES | 38.5 |
| CAP1_7 | NALFNSIC(K) | 9 | 2 or 3 | 2 | 0 | 0.046 | YES | 26.38 |
| CAP1_8 | TGPVASAGAAPP(R) | 13 | 2 | 4 | 0 | 0.096 | YES | 13.44 |
| GCN1_3 | LTSPEALRPSVVNITGPLI(R) | 20 | 3 | 2 | 0 | 0.049 | YES | 39.42 |
| GCN1_4 | FVIQGAGS(K) | 9 | 2 | 2 | 0 | - | YES | 18.47 |
| GCN1_5 | IIIDDLLEAT( R) | 11 | 2 | 2 | 0 | 0.045 | YES | 39.06 |
| FBP1_1 | GTGELTTLLNAVGTAV(K) | 17 | 3 | 4 | 0 | 0.031 | YES | 39.25 |
| FBP1_2 | SSFTSCVLVSEEND(K) | 15 | 3 | 2 | 0 | 0.494 | YES | 27.6 |
| UQCRFS1_2 | LTDIPEG(K) | 8 | 2 | 2 | 0 | - | YES | 15.11 |
| UQCRFS1_2 | TVVSQFISSMSASADVLALS(K) | 21 | 3 or 4 | 2 | 0 | 0.037 | YES | 45.21 |
| UQCRFS1_3 | EIATEEAVNLAELRDPQHDKD(R) | 22 | 4 | - | 0 | - | YES | 32.46 |
| GHITM_1 | AETHPLYGVQ(K) | 11 | 3 | 2 | 0 | - | YES | 18.79 |
| GHITM_2 | EAAFEPATDTAI( R) | 13 | 2 or 3 | 4 | 0 | 0.049 | YES | 25.13 |
| GHITM_3 | SVIWPQYV(K) | 9 | 2 | 5 | 0 | - | YES | 32.65 |
